# The force-sensing GPCR LPHN2 is indispensable for normal auditory function

**DOI:** 10.1101/2023.10.24.563883

**Authors:** Zhao Yang, Ming-Wei Wang, Shu-Hua Zhou, Zhi-Chen Song, Kong-Kai Zhu, Xin Wen, Qi-Yue Zhang, Ying Guan, Jia-Rui Gao, Xiao-Hui Wang, Ya-Qi Wang, Wen-Wen Liu, Lei Xu, Wei Xiong, Ren-Jie Chai, Chuan Wang, Zhi-Gang Xu, Xiao Yu, Jin-Peng Sun

## Abstract

The conversion of force sensation into electrical signals via mechanoelectrical transduction (MET) is considered the key step in auditory perception. Here, we found that G protein-coupled receptor (GPCR) LPHN2/ADGRL2 was expressed at the tips of stereocilia in cochlear hair cells and was associated with MET channel components. Hair cell-specific LPHN2 deficiency caused hearing loss and impaired MET responses. A specific inhibitor of LPHN2, developed by in silico screening and pharmacological characterization, reversibly blocked the MET response. Mechanistically, administration of force to LPHN2 activated TMC1 through direct interaction and caused conformational changes in TMC1 in vitro. Furthermore, the sensing of force by LPHN2 stimulated Ca^2+^ responses and neurotransmitter release in hair cells. Finally, hearing loss in LPHN2-deficient mice was reversed by the re-expression of LPHN2-GAIN in cochlear hair cells. The important roles of LPHN2 in auditory perception and a TMC-coupled force sensor indicated that GPCRs could be candidate auditory receptors.

## Introduction

Auditory perception is one of our basic senses and allows us to experience the features of the acoustic world, including melodious music, chirps of birds and tinkling of spring water. Additionally, auditory perception is our main mechanism for communicating with each other via speech, and it alerts us to possible dangers. By converting mechanical force into electrical signals, previous studies have suggested that mechanoelectrical transduction (MET) in cochlear hair cells act as the central event during auditory perception (Gillespie and Muller, 2009; Hudspeth and Corey, 1977; Hudspeth and Jacobs, 1979; Kawashima et al., 2011; Pan et al., 2018; Zheng and Holt, 2021). The molecular identity of MET channels in hair cells remained elusive for decades; however, a series of candidate components of the MET channel complex, which include the transmembrane membrane proteins TMC1/2, TMIE, and LHFPL5 as well as auxiliary subunits such as calcium (Ca^2+^) and integrin binding protein 2 and 3 (CIB2/3), were recently identified and have gradually been characterized (Cunningham et al., 2020; Giese et al., 2017; Liang et al., 2021; Pan et al., 2013; Tang et al., 2020; Wang et al., 2023; Xiong et al., 2012). In the research field of hearing, TMC1/2 are increasingly becoming recognized as the pore-forming subunits of hair cell MET channels. In addition to these MET components, which are concentrated at the tips of stereocilia, another mechanosensitive ion channel Piezo2, which is expressed at the apical surface of cochlear hair cells, has also been suggested to be a force sensor that is required for reverse-polarity currents (Wu et al., 2017).

In addition to ion channels, G protein-coupled receptors (GPCRs) are well known to function as important sensory receptors that mediate the sensation of light, odor, itch and at least three qualities of taste (Buck and Axel, 1991; Guo et al., 2023; Yang et al., 2021; Yau and Hardie, 2009; Zhang et al., 2003). Importantly, several GPCRs have been shown to be essential for hearing. For instance, CELSR1 and Fzd3/6 are required for the maintenance of cochlear planar cell polarity, and LGR5 regulates the self-renewal abilities of cochlear hair cells (Chai et al., 2012; Duncan et al., 2017; Shi et al., 2013; Wang et al., 2006). Moreover, ADGRV1 is a component of ankle links and is required for proper stereocilia development (Guan et al., 2023; Hu et al., 2014; McGee et al., 2006; Weston et al., 2004). Although GPCRs play important regulatory roles in cochlear development, it is still unknown whether GPCRs participate in direct force sensation during hearing processes, modulate membrane excitability and contribute to neurotransmitter release. In contrast to ion channels, GPCRs often modulate intracellular secondary messengers to produce sensory signals (Arshavsky and Bownds, 1992; Chandrashekar et al., 2006; Chen and Yau, 1994; Nakamura and Gold, 1987; Yang et al., 2021; Zhang et al., 2003). Notably, the seminal work of the Hudspeth Laboratory showed that MET is an extremely rapid process that occurs within microseconds (Corey and Hudspeth, 1979). Therefore, GPCRs are generally not considered to be direct auditory receptors because of their slower kinetics. For example, the latency of the fastest known G protein signaling pathway, which is mediated by the rhodopsin-Gt-TRP channel complex and participates in Drosophila phototransduction, was measured to be ∼20 ms.

In addition to the conventional secondary messenger system, we hypothesized that GPCRs may also be able to regulate acute physiological responses by directly binding to and modulating the activity of ion channels via conformational changes (Davare et al., 2001; Liu et al., 2017; Zhang et al., 2012). Therefore, it is possible that GPCRs could serve as hearing sensors and may interact with the quick initial sensation of sound that is mediated by mechanosensitive ion channels or individually trigger neurotransmitter release. We hypothesized that candidate auditory receptors (exemplified by GPCRs) may meet the following criteria: (1) the candidate is expressed in the stereocilia of cochlear hair cells; (2) the candidate directly senses force in a physiological range (2∼100 dynes/cm^2^); (3) the candidate can convert force stimuli to modulate the membrane potentials of hair cells, mediate changes in intracellular second messengers or regulate neurotransmitter release; and (4) knocking out the gene that encodes the candidate leads to hearing disorders in animal models. Considering these criteria, we cannot exclude the possibility that GPCRs might directly participate in auditory perception by physically interacting with MET subunits or modulating specific variants, such as amplitude or period, during hearing sensation.

In our parallel paper, we found that a mechanosensitive GPCR, LPHN2, was uniquely expressed at the apical surface of utricular hair cell, which was indispensable for normal equilibrioception of mice (Yang et al., 2024). In the present study, we showed that LPHN2 was distributed in cochlear hair cells at both the stereocilia and the apical membrane surface. Global knockout of *Lphn2* or inducible hair cell-specific LPHN2 knockout impaired the hearing of mice. Importantly, the MET current in LPHN2-expressing cochlear hair cells was impaired either by genetic ablation of LPHN2 or by pharmacological inhibition of LPHN2 with a specific inhibitor. Mechanistically, when LPHN2 sensed force, it directly coupled to TMC1 and induced conformational changes in TMC in an *in vitro* reconstitution system, and these effects occurred independently of its Gs-activating ability. We further revealed that force sensation by LPHN2 induced Ca^2+^ signaling and transmitter release in cochlear hair cells. Overall, our findings identify a mechanosensitive GPCR that could function as a direct sensor of force during hearing by functionally coupling to TMC1; this GPCR might be a component of the mechanotransduction machinery of cochlear hair cells.

## Results

### Expression of LPHN2 in cochlear hair cells

Previous single-cell RNA sequencing (scRNA-seq) data revealed that LPHN2 is expressed primarily in cochlear hair cells and that its expression exhibits a dynamic pattern (Ranum et al., 2019). Our quantitative RT-PCR results revealed that the mRNA level of *Lphn2* in the mouse cochlear epithelium decreased by approximately 30% from E20 to P14 and then reached a steady-state at P30 (Figure 1A). Using the purified LPHN2 protein as a reference, the protein level of LPHN2 in cochlear hair cells was measured to be approximately 387 ± 13 fmol/mg, which is approximately 1.5-fold higher than that of VLGR1, at P10 (Figures S1A and S1B). VLGR1 is a deafness-associated GPCR and an ankle link component of hair cells (Guan et al., 2023). To determine the pattern of LPHN2 expression in the cochlea, we performed whole-mount immunostaining with a commercial anti-LPHN2 antibody; the specificity of this antibody was demonstrated in both HEK293 cells overexpressing LPHN2 and cochleae derived from *Lphn2^-/-^* mice at E18 (Figures S1C and S1D). LPHN2 expression was detected in approximately 80% of both outer hair cells (OHCs) and inner hair cells (IHCs) throughout the length of the cochlear duct at P10, and there were no significant tonotopic differences in the apical, middle or basal turns (Figure 1B).

**Figure 1.**
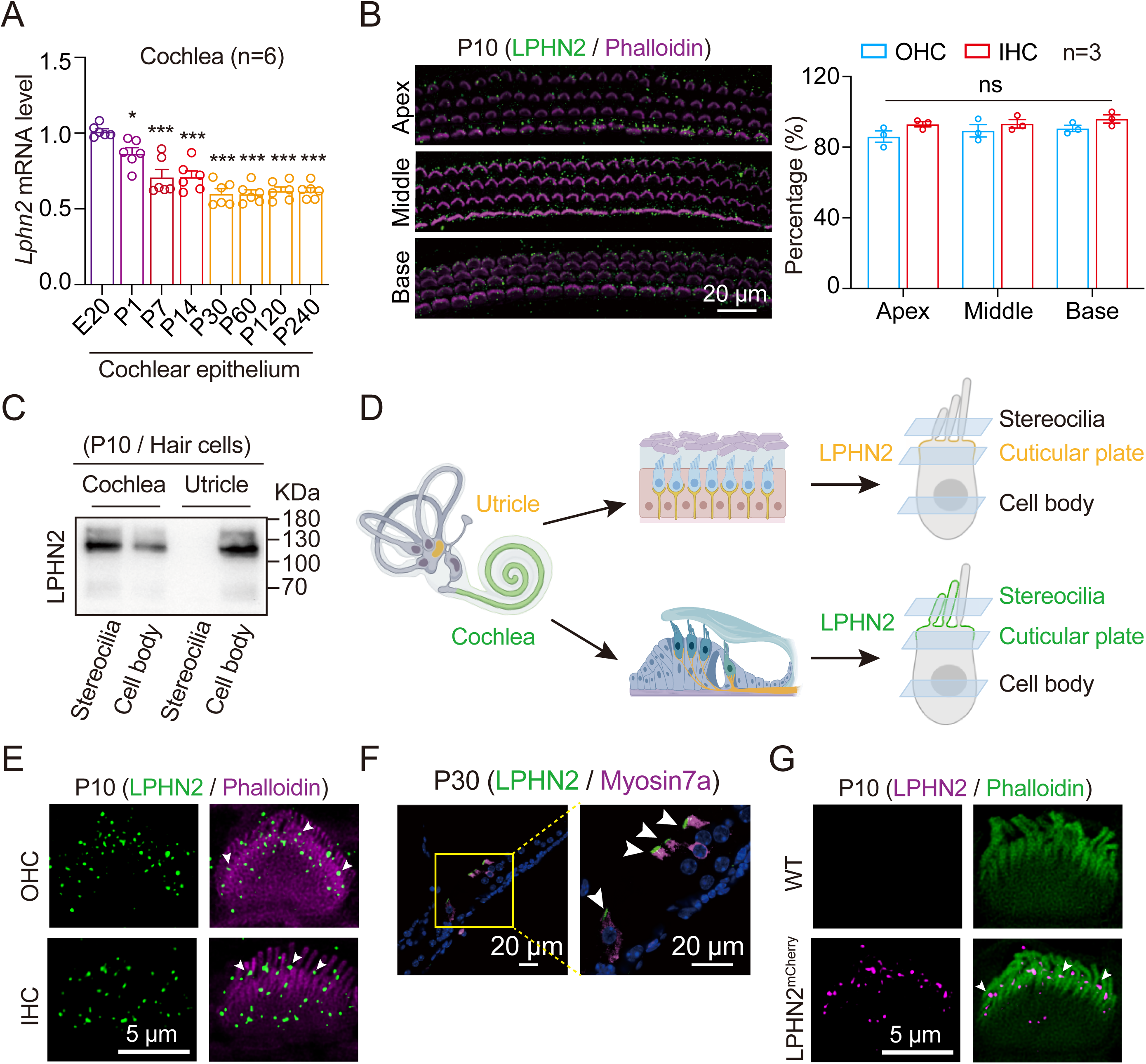
Expression of LPHN2 in cochlear hair cells. **(A)** mRNA levels of *Lphn2* in the cochlear epithelium isolated from WT mice at E20, P1, P7, P14, P30, P60, P120 and P240 (n = 6). Data are normalized to the mRNA levels of *Lphn2* at E20. E, embryonic period; P, postnatal day. **(B)** Immunostaining (left) and quantitative analysis (right) of LPHN2 expression in hair cells from different sections of cochlear whole mounts derived from WT mice at P10 (n = 3 mice per group). Scale bar, 20 μm. **(C)** Representative blotting showing the protein levels of LPHN2 in the stereocilia and cell body of cochlear or utricle hair cells isolated from WT mice at P10. Representative blotting from three independent experiments are shown (n = 3). LPHN2 antibody against the N terminus (551 - 600 residues) was used, which revealed both the full-length (∼130 kDa) and the N-terminal fragment of LPHN2 (∼120 kDa) in the cochlear or utricle hair cells. Data are correlated to Figure S1F-S1G. **(D)** Schematic diagram showing the expression pattern of LPHN2 in cochlear and utricle hair cells, respectively. **(E)** Co-immunostaining of LPHN2 (green) and Phalloidin (magenta) in the stereocilia of OHC top) and IHC (bottom) in cochlear whole mounts derived from WT mice at P10 (n = 6 mice per group). Scale bar, 5 μm. White arrows show the localization of LPHN2 at the tips of shorter rows of stereocilia. Data are correlated to Figure S1H. **(F)** Co-immunostaining of LPHN2 (green) and Myosin7a (magenta) in cochlear sections derived from WT mice at P30 (n = 6 mice per group). Scale bar, 20 μm. White arrows show the distribution of LPHN2 above the apical surface of hair cells. **(G)** Expression of LPHN2-mCherry (magenta) in the stereocilia of IHC in cochlear whole mounts derived from WT or LPHN2^mCherry^ mice at P10 (n = 6 mice per group). Scale bar, 5 μm. White arrows show the localization of LPHN2 at the tips of shorter rows of stereocilia. Data are correlated to Figure S1I-S1J. **(A)** *P < 0.05; ***P < 0.001. WT mice at postnatal time points compared with those at E20. **(B)** ns, no significant difference. Hair cells at middle or basal turn compared with those at apical turn. The bars indicate mean ± SEM values. All data were statistically analyzed using one-way ANOVA with Dunnett’s post hoc test.

Importantly, via separation of stereocilia from the hair cell body using a modified twist-off method (Avenarius et al., 2017), we were able to detect LPHN2 expression in both the stereocilia and the cell bodies of cochlear hair cells using TMIE/PCDH15 and Myosin7a as markers of stereocilia and cell bodies, respectively (Figures 1C and S1E-S1G). The results showed that LPHN2 was expressed only in the cell body but not in the stereocilia of utricular hair cells, which was consistent with the immunostaining results of our parallel study (Figures 1C and S1E-S1G). To further explore the subcellular expression pattern of LPHN2, we collected different optical sections ranging from the stereocilia to the cochlear hair cell body and examined the localization of LPHN2 (Figure 1D). Notably, in both the cochlear OHCs and IHCs of mice at P10, immunofluorescence puncta indicating LPHN2 were observed along the stereocilia, with staining at the tips of several shorter rows of stereocilia but not in the longest stereocilia (Figure 1E). LPHN2 immunofluorescence was also observed at the apical surfaces above the cuticular plate of HCs; this conclusion was supported by the close proximity of LPHN2 to spectrin, which is a specific marker of the cuticular plate, and by the results of staining sections of the cochlear sensory epithelium with an antibody against Myosin7a, which is a marker of hair cells (Figures 1F and S1H). In contrast, LPHN2 was not expressed at the basolateral membrane along the cell bodies of HCs (Figure 1F). To exclude the possibility of antibody nonspecificity, we detected LPHN2 expression in whole mounts of cochlea derived from a transgenic knock-in mouse line in which the C-terminus of LPHN2 is tagged with an mCherry sequence (LPHN2^mCherry^ mice) (Figure S1I). Consistent with the immunostaining data, LPHN2-mCherry was also expressed at the tips of shorter rows of stereocilia as well as at the apical surface of both IHCs and OHCs (Figures 1G and S1J). The specific expression pattern of the mechanosensitive receptor LPHN2 in cochlear hair cells, especially its localization at the tips of stereocilia, suggests that LPHN2 might perform unique functions in the auditory system.

### LPHN2 deficiency severely impairs hearing

To investigate the functional roles of LPHN2 in hearing, we initially used heterozygous *Lphn2* gene-knockout (*Lphn2*^+/-^) mice since homozygous knockout of this gene (*Lphn2*^-/-^ mice) results in embryonic lethality (Anderson et al., 2017) (Figures S2A-S2C). *Lphn2*^+/-^ mice are viable and exhibit normal growth, but vestibular defects were observed in these mice in our parallel study (Yang et al., 2024). Measurement of the auditory brainstem response (ABR) revealed that at P30, *Lphn2*^+/-^ mice had an ∼30 dB higher auditory threshold in response to click stimuli than did their wild-type (WT) littermates (Figure S2D). Moreover, compared with those of WT mice, *Lphn2*^+/-^ mice exhibited significantly elevated ABR thresholds at different frequencies (4, 8, 12, 16, 24 and 32 kHz) (Figures S2D and S2E). ABR wave I analysis further indicated a decreased wave amplitude and increased wave latency in *Lphn2*^+/-^ mice, suggesting that the function of IHC synapses was impaired in *Lphn2*^+/-^ mice (Figure S2F). Furthermore, the threshold of distortion product otoacoustic emission (DPOAE) in *Lphn2*^+/-^ mice was significantly increased between 12 and 24 kHz, suggesting that OHC function was affected in *Lphn2*^+/-^ mice (Figure S2G). Since the deterioration of hearing function might occur due to developmental defects in hair cells, we examined hair cell morphology by immunostaining and scanning electron microscopy. The results of both analyses revealed no detectable impairment in the structure or morphology of OHCs or IHCs in the cochleae of *Lphn2*^+/-^ mice, suggesting that the heterozygous ablation of LPHN2 in mice did not cause developmental defects in cochlear hair cells, at least at the age of 4 weeks (Figures S2H-S2J).

To further assess the specific role of LPHN2 in cochlear hair cells, we crossed *Lphn2^fl/fl^* mice with an inducible *Pou4f3-CreER^+/-^*transgenic mouse line (Figure S2K). Previous scRNA-seq data (GSE114157) indicated that POU4F3, which is a commonly used marker of hair cells, is expressed in all LPHN2-positive cochlear hair cells; therefore, we used *Pou4f3-CreER^+/-^ Lphn2^fl/fl^* mice to knockout LPHN2 specifically in hair cells. Consistently, the specific knockout of LPHN2 in the cochleae and utricles of *Pou4f3-CreER^+/-^Lphn2^fl/fl^* mice was confirmed by both immunofluorescence staining and western blotting (Figures S2L-S2N). In contrast, LPHN2 expression in other tissues, such as the brain, was not significantly affected in *Pou4f3-CreER^+/-^ Lphn2^fl/fl^* mice (Figures S2L and S2M). Notably, compared with their control *Pou4f3-CreER^+/-^ Lphn2^+/+^* littermates, *Pou4f3-CreER^+/-^Lphn2^fl/fl^* mice exhibited severe hearing impairment, as indicated by an elevated ABR threshold at all testing frequencies and deterioration of DPOAE between 8 and 32 kHz (Figures 2A-2D). As in *Lphn2*^+/-^ mice, *Pou4f3-CreER^+/-^Lphn2^fl/fl^* mice exhibited normal cochlear hair cell structure and morphology at P30 (Figures 2E, 2F and S2O). In addition, the expression levels and the localization of MET machinery components, including TMC1, TMC2, TMIE, LHFPL5 and PCDH15, in the cochlear hair cells of *Pou4f3-CreER^+/-^Lphn2^fl/fl^* mice were comparable to those in their wild-type littermates (Figure 2G). Collectively, these findings support the idea that LPHN2 expressed in cochlear hair cells is indispensable for normal hearing.

**Figure 2.**
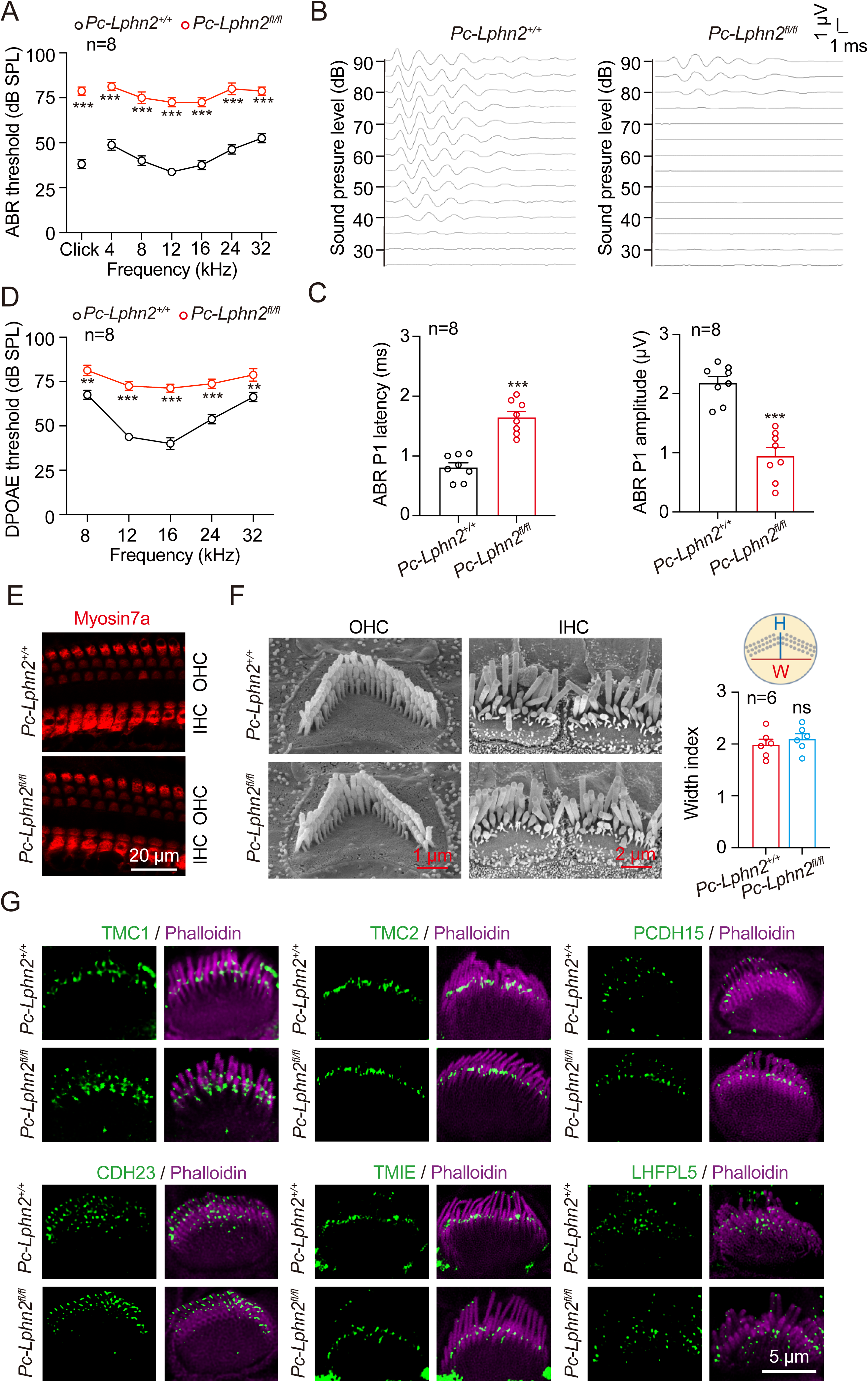
LPHN2 deficiency severely impairs hearing. **(A-D)** ABR thresholds (A), ABR waveforms at click (B), ABR peak1 (P1) latency (left) and amplitude (right) (C), and DPOAE thresholds (D) of *Pou4f3-CreER^+/-^Lphn2^+/+^* (referred to as *Pc-Lphn2^+/+^*) and *Pou4f3-CreER^+/-^Lphn2^fl/fl^* (referred to as *Pc-Lphn2^fl/fl^*) mice at P30 (n = 8 mice per group). **(E)** Immunostaining of Myosin7a (red) in cochlear hair cells derived from *Pc-Lphn2^+/+^* and *Pc-Lphn2^fl/fl^* mice at P30 (n = 6 mice per group). Scale bar, 20 μm. Data are correlated to Figure S2O. **(F)** Left: Representative SEM images showing hair bundle morphology of the OHC and IHC from *Pc-Lphn2^+/+^* and *Pc-Lphn2^fl/fl^* mice at P30. Scale bar, 1 μm (OHC) or 2 μm (IHC). Right: Quantitative analysis of the width index of hair bundles of the OHC from *Pc-Lphn2^+/+^* and *Pc-Lphn2^fl/fl^* mice at P30 (n = 6 mice per group). **(G)** Co-immunostaining of Phalloidin (magenta) and different MET machinery components (green), including TMC1, TMC2, PCDH15, CDH23, TMIE and LHFPL5, in cochlear hair cells of *Pc-Lphn2^fl/fl^* and *Pc-Lphn2^+/+^* mice (n = 6 mice per group). **(A, C-D, F)** **P < 0.01; ***P < 0.001; ns, no significant difference. *Pc-Lphn2^fl/fl^* mice compared with *Pc-Lphn2^+/+^* mice. The bars indicate mean ± SEM values. All data were statistically analyzed using two-way ANOVA with Dunnett’s post hoc test (A, D) or unpaired two-sided student’s *t* test (C, F).

### Force sensation by LPHN2 induces changes in second messengers in cochlear hair cells

GPCRs act as direct sensors of itch and participate in the senses of vision, smell, and taste by converting extracellular stimuli into changes in the intracellular levels of second messengers, such as cAMP, cGMP and Ca^2+^ (Filipek et al., 2003; Guo et al., 2023; Kaupp, 2010; Smith, 2023; Yang et al., 2021; Yang et al., 2023). Our parallel study indicated that LPHN2 activated Gs and Gi signaling in response to force stimulation (Figures S3A and S3B). We therefore measured cAMP levels after the delivery of force to LPHN2 using paramagnetic beads that were coated with an anti-LPHN2 antibody (LPHN2-M-beads). This antibody recognizes the N-terminus (residues 551-600) of LPHN2 and can exert quantifiable force on LPHN2 in a magnetic field (Figure 3A). In the heterologous system, we showed that upon the application of magnetic force, LPHN2-M-beads induced a dose-dependent increase in the cAMP concentration in LPHN2-overexpressing HEK293 cells but not in cells that were transfected with the empty vector pcDNA3.1 (Figures 3B and S3C). The EC50 for the LPHN2-M-bead-induced increase in cAMP concentrations in vitro was 1.70 ± 0.34 pN. In contrast, polylysine-coated paramagnetic beads (Ctrl-beads) did not increase the cAMP concentrations in LPHN2-expressing HEK293 cells in response to the application of force, thus supporting the specificity of the LPHN2-M-beads in activating LPHN2 (Figure 3B). We subsequently applied LPHN2-M-beads to murine cochlear explants. The results revealed that the application of 3 pN force to cochlear sensory epithelium harvested from *Pou4f3-CreER^+/-^Lphn2^+/+^* mice induced an approximately 1.6∼2.2-fold increase in intracellular cAMP levels in different sections (apical turn: from 1.49 ± 0.11 mM to 2.41 ± 0.13 mM; middle turn: from 1.58 ± 0.07 mM to 3.41 ± 0.18 mM; basal turn: from 1.51 ± 0.08 mM to 2.48 ± 0.14 mM); these effects of force were suppressed in cochlear explants that were pretreated with the nonspecific Gs inhibitor NF449 or in cochlear explants from *Pou4f3-CreER^+/-^Lphn2^fl/fl^* mice at P10 (Figure 3C). In contrast, the force applied by Ctrl-beads did not induce detectable changes in cAMP concentrations in cochlear explants (Figure S3D). These results suggest that the Gs-cAMP pathway downstream of LPHN2 is activated in response to force stimulation in the cochlea. Consistent with these findings, Gs was the most abundantly expressed G protein subtype in LPHN2-expressing cochlear hair cells, as revealed by both scRNA-seq data and single-cell qRT‒PCR analysis of 20 isolated cochlear hair cells (Figures 3D and S3E).

**Figure 3.**
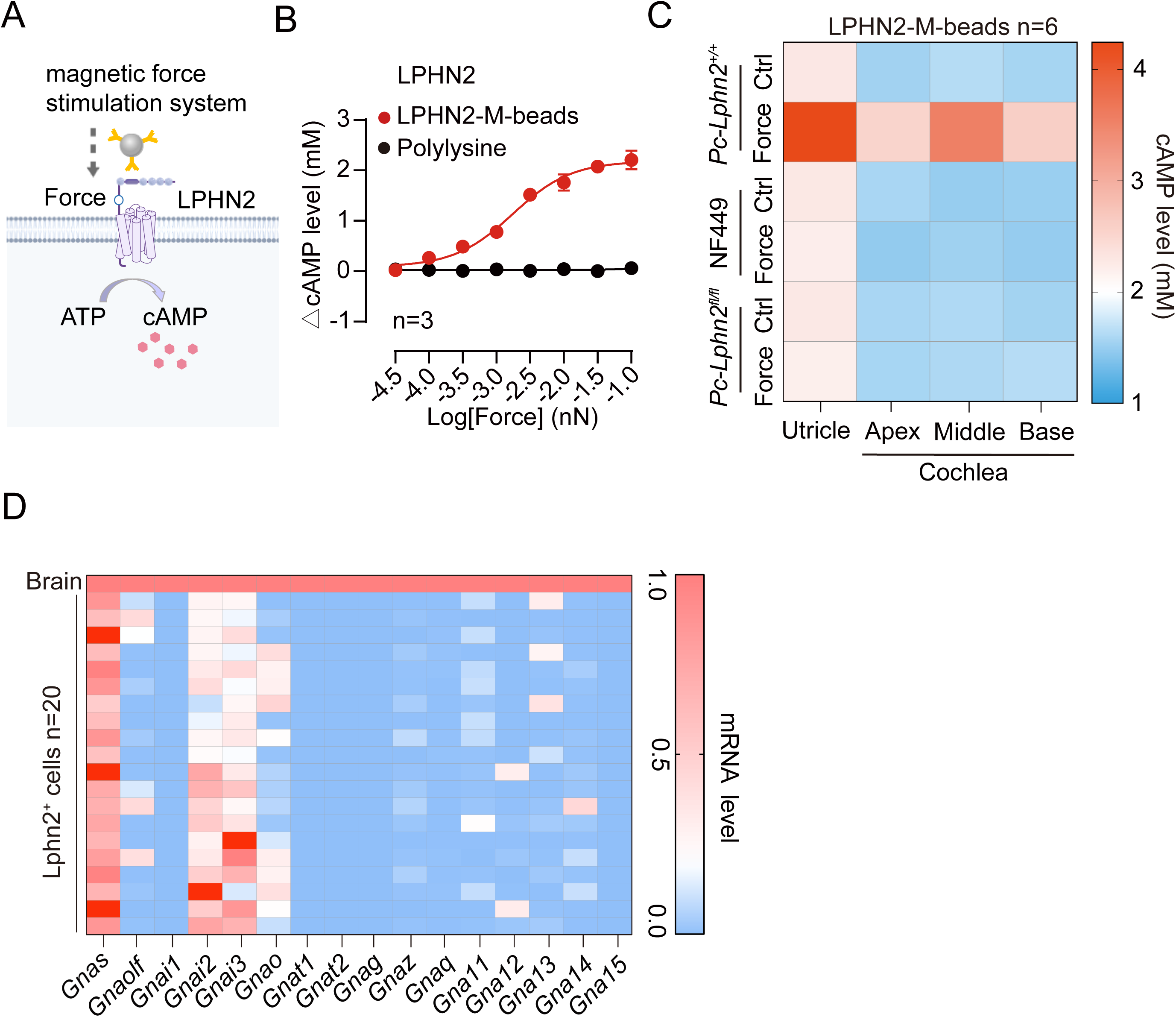
Force sensation by LPHN2 induces endogenous cAMP production. **(A)** Schematic representation of the force-induced LPHN2 activation in HEK293 cells using a magnetic system. **(B)** Dose-dependent cAMP accumulation in HEK293 cells transfected with plasmid encoding LPHN2 in response to force applied via LPHN2-M-beads or polylysine-coated Ctrl-beads (n = 3). Data are correlated to Figure S3C. **(C)** Endogenous cAMP level changes in the cochlea (apex, middle or base) and utricle explant derived from *Pc-Lphn2^+/+^*or *Pc-Lphn2^fl/fl^* mice at P10 in response to force (3 pN) applied by LPHN2-M-beads in the absence or presence of NF449 (20 μM) (n = 6). Data are correlated to Figure S3D. **(D)** mRNA levels of different G protein subtypes in LPHN2-expressing cochlear hair cells derived from P10 mice measured by single-cell qRT-PCR (n = 20 hair cells). The mRNA levels of respective G protein subtypes in the brain were used as the reference (normalized to 1).

Notably, the intensity of the force that was applied by magnetic beads in this study was determined based on our calculation that 10 pN equals 100 dynes/cm^2^ on the plasma membrane of stereocilia or the apical surface of hair cells; this approach was used in a previous study to mimic arterial wall shear stress (approximately 1 m/s under physiological conditions) (Mehta et al., 2020). Therefore, the force of 1.7∼10 pN that was used in the present study was within the physiological range, mimicking approximately 17 cm/s ∼100 cm/s fluid velocity.

### LPHN2 deficiency impairs the MET current in cochlear hair cells

The localization of LPHN2 at the tips of shorter rows of stereocilia in cochlear hair cells, the observation of auditory defects in *Lphn2*-deficient mice, and the lack of abnormalities in hair bundle development or MET component assembly in *Lphn2*-deficient mice collectively suggest that LPHN2 might directly participate in auditory perception. We therefore investigated whether LPHN2 participates in MET by performing whole-cell voltage-clamp recording experiments. In these experiments compared recordings of mid-apical cochlear hair cells derived from *Pou4f3-CreER^+/-^Lphn2^fl/fl^* mice with those of cells derived from wild-type littermate mice in response to deflection of hair bundles by a fluid jet (Figure 4A). To record MET currents specifically in LPHN2-expressing hair cells, we administered a modified AAV-ie-*Lphn2pr*-mCherry to P3 mice via round window membrane injection (Figure 4A). The efficiency of mCherry labeling and accuracy of viral targeting of LPHN2-expressing cells were validated by both immunostaining with an anti-LPHN2 antibody and single-cell RT‒PCR (Figures S4A and S4B).

**Figure 4.**
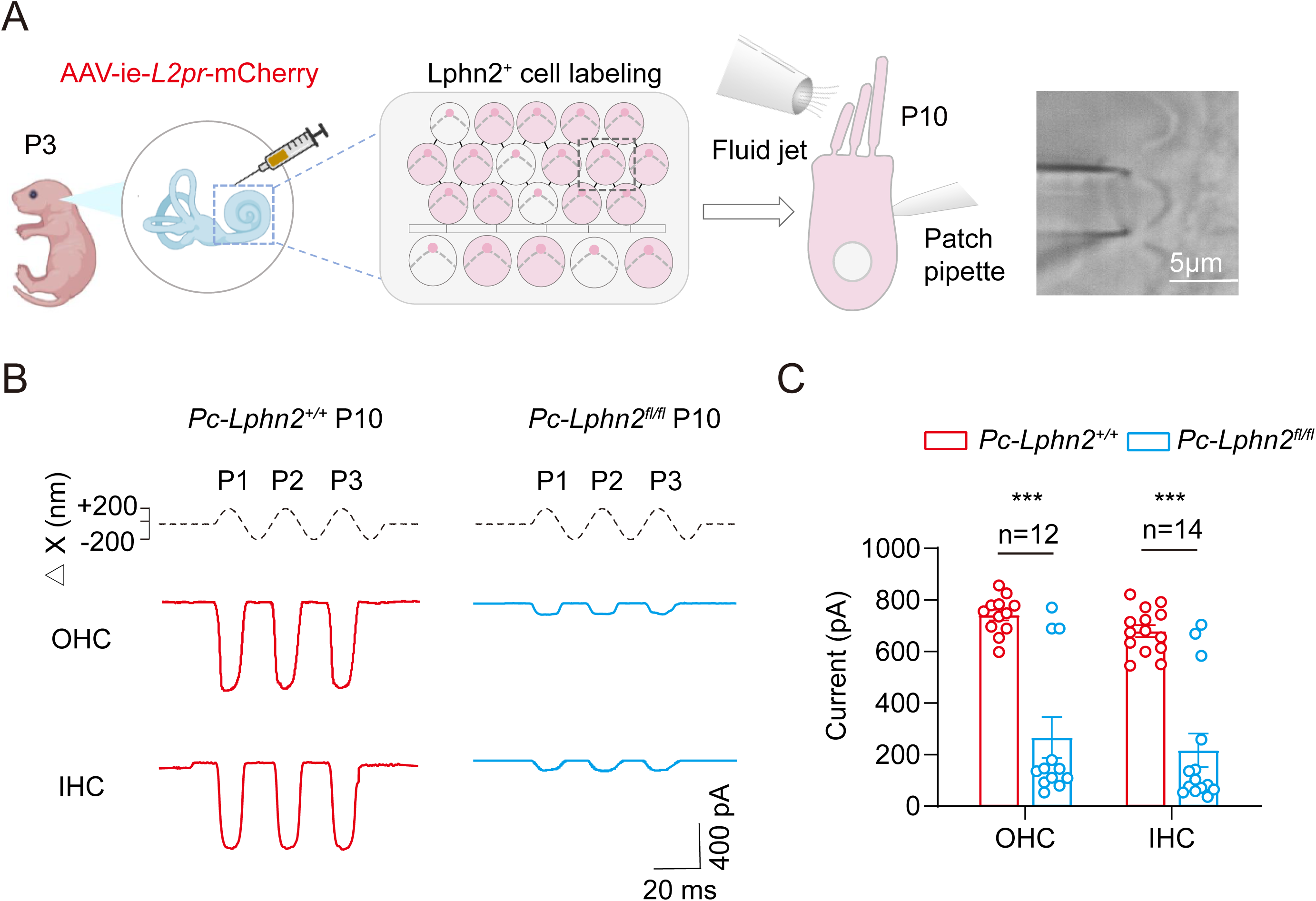
LPHN2 deficiency impairs MET currents. **(A)** Schematic illustration of the labelling of LPHN2-expressing hair cells by AAV-ie-*Lphn2pr*-mCherry (referred to as AAV-ie-*L2pr*-mCherry) and the MET current recording by fluid-jet stimulation. Data are correlated to Figure S4A-S4B. **(B)** Representative MET current traces induced by sinusoidal fluid-jet stimulation in cochlear OHCs (top) and IHCs (bottom) of *Pc-Lphn2^+/+^* mice (red) or *Pc-Lphn2^fl/fl^* mice (blue) at P10. **(C)** Quantification of the saturated MET currents in cochlear OHCs and IHC of *Pc-Lphn2^+/+^* mice or *Pc-Lphn2^fl/fl^* mice at P10 (n = 12 for OHCs and n = 14 for IHCs). **(C)** ***P < 0.001; *Pc-Lphn2^+/+^* mice compared with *Pc-Lphn2^fl/fl^* mice. The bars indicate mean ± SEM values. All data were statistically analysed using unpaired two-sided Student’s *t* test.

The cochleae of AAV-treated mice were isolated at P10, and mCherry-labeled hair cells were selected to record macroscopic MET currents at a holding potential of -80 mV. In the LPHN2-expressing OHCs from control *Pou4f3-CreER^+/-^Lphn2^+/+^* mice, the excitatory stimuli at maximal deflection elicited peak MET currents of 741.6 ± 25.4 pA (Figures 4B and 4C). In contrast, compared with those of control littermates, the peak MET currents in the mCherry-labeled OHCs from *Pou4f3-CreER^+/-^Lphn2^fl/fl^*mice were reduced by approximately 70% (to 235.9 ± 83.3 pA) (Figures 4B and 4C). Similar defective MET currents were also observed in LPHN2-expressing IHCs from *Pou4f3-CreER^+/-^Lphn2^fl/fl^*mice (*Pou4f3-CreER^+/-^Lphn2^fl/fl^* 216.4± 65.1 pA vs. *Pou4f3-CreER^+/-^Lphn2^+/+^* 679.7 ± 23.1 pA) (Figures 4B and 4C). These results indicated that LPHN2 plays important roles in normal mechanotransduction in cochlear hair cells.

### Development of a selective LPHN2 inhibitor that antagonizes its force sensation

In addition to genetic LPHN2 knockout, a reversible and selective inhibitor of LPHN2 that antagonizes its force sensation could be a very useful tool for investigating how LPHN2 participates in hearing sensation and MET, and such an inhibitor could exclude the possibility of developmental defects caused by genetic ablation. We therefore constructed an inactive LPHN2 structural model according to the inactive CD97 structure that we recently solved, and we rationally designed an in-silico screening strategy to search for LPHN2 inhibitors (Figure 5A) (Mao et al., 2024a). A total of 1000,000 compounds with diverse skeleton types in a designed compound database were docked into the predicted top binding pocket of inactive LPHN2 to identify the top 5000 small molecules according to docking scores (Figure 5B). To achieve high selectivity, we filtered 5000 compounds according to their interactions with specific residues (I855, E894, F921, L1055, and T1070) in the LPHN2 pocket, because these five residues are not well conserved among aGPCR families (Figure S5A). After performing cluster analysis of the top 300 small molecules, we identified 48 compounds that exhibited high structural diversity (Figure 5B). We then performed the BRET-based G protein dissociation assay with LPHN2-overexpressing HEK293 cells, and we determined that one of these compounds, namely D11, inhibited 3 pN force-induced LPHN2 activation in a concentration-dependent manner, with an EC50 of 39.88 ± 16.24 nM (Figures 5C-5E). To further predict the binding mode of D11 to LPHN2, we performed computational simulation using a Ligand Docking panel inserted in Maestro. D11 was modeled in the orthosteric pocket of LPHN2 and exhibited a slanted V-shaped configuration (Figure 5F). Twelve hydrophobic residues (L890^2.57^, F897^2.64^, I901^ECL1^, L917^3.36^, F921^3.40^, F925^3.44^, W983^ECL2^, L1047^6.49^, W1051^6.53^, L1055^6.57^, F1057^ECL3^, and F1069^7.42^) and three polar residues (E894^2.61^, N1073^7.46^ and Q1076^7.49^) from TM2-TM3, TM6-TM7 and ECL1-ECL3 of LPHN2 formed the potential D11 binding pocket (Figures 5F and 5G). In particular, the carboxyl oxygen in thiazide ring of D11 formed polar interactions with the side chain oxygen atom of E894^2.61^ and side chain nitrogen atom of N1073^7.46^ (Figure 5G). The trifluoromethylbenzene ring of D11 formed π-π interactions with F897^2.64^ and W983^ECL2^ of LPHN2 (Figure 5G). Besides, the trifluoromethylbenzene ring of D11 formed hydrophobic interactions with I901^ECL1^, L917^3.36^, L1055^6.57^, F1057^ECL3^, and F1069^7.42^. The 1,4-dimethoxyphenyl group inserted in a hydrophobic pocket composed by L890^2.57^, F921^3.40^, F925^3.44^, L1047^6.49^ and W1051^6.53^. Consistently, mutations in residues of the D11 binding pocket, such as E894^2.61^A, F921^3.40^A, W983^ECL2^A or F1069^7.42^A, but not mutations in other randomly selected surrounding residues, abolished the inhibitory effects of D11 on force-induced LPHN2 activation (Figures 5H, S5B and S5C). Importantly, D11 inhibited only force-induced LPHN2 activation, and it did not have detectable effects on either the force-induced or agonist-induced activation of other adhesion GPCRs in the heterologous HEK293 overexpression system (Figure 5I). These results suggested that D11 is a selective LPHN2 inhibitor. Overall, we identified a specific blocker of force sensation by LPHN2 through a rationally designed in silico screening method.

**Figure 5.**
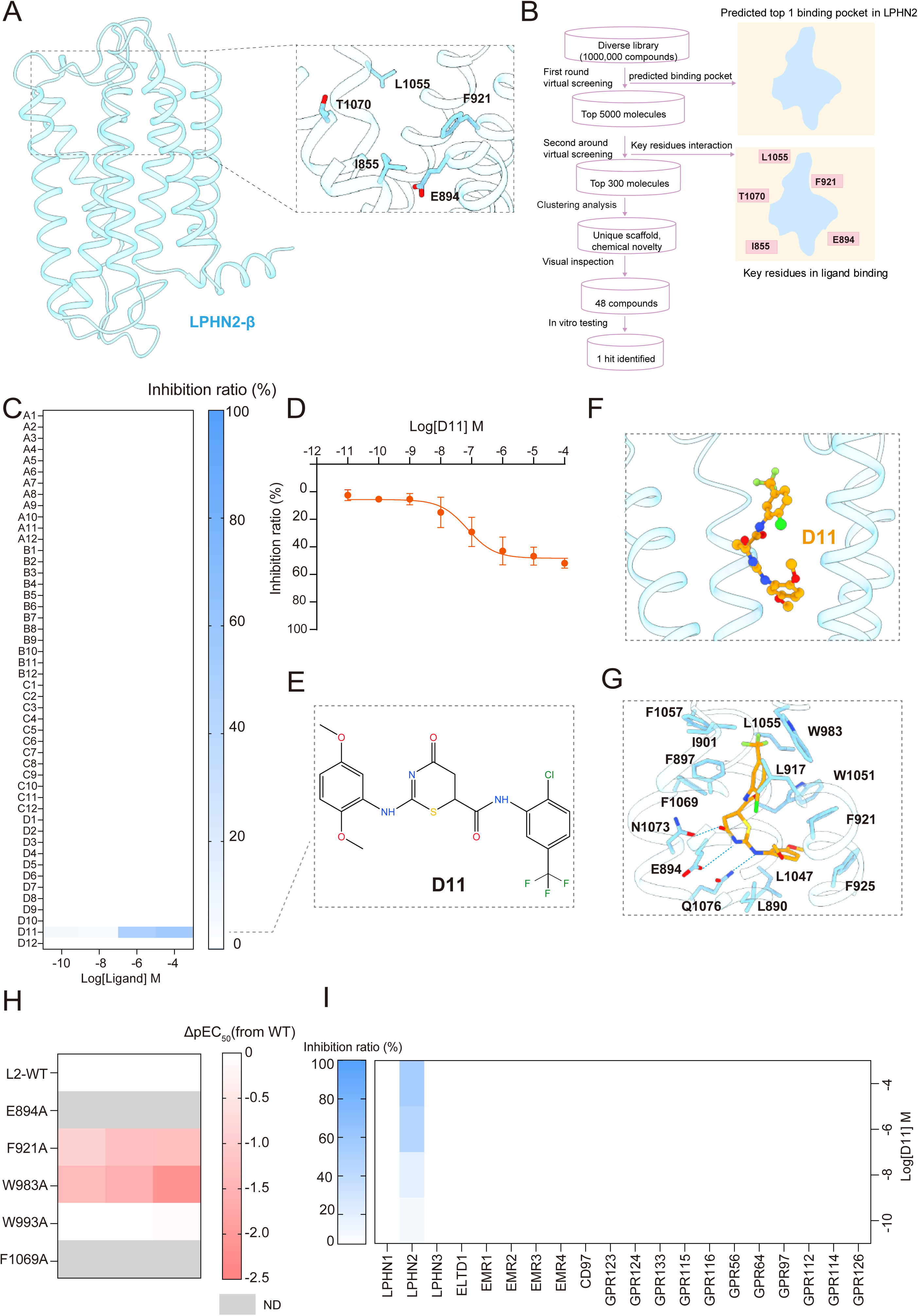
Identification of selective LPHN2 inhibitor antagonizing its force sensation. **(A)** Simulated inactive LPHN2-β structural model according to the cryo-EM structure of the inactive CD97 (PDB: 8IKJ). The LPHN2-specific pocket residues that are not conserved among aGPCRs are shown. **(B)** The procedures for in-silico screening of selective LPHN2 inhibitors. 100 million compounds were virtually screened against the putative orthosteric pocket of LPHN2 and the top 5000 were selected for the second round of screening against the LPHN2-specific pocket residues. The resultant top 300 molecules were subjected to clustering analysis, leading to 48 candidates. The final hit was identified by *in vitro* BRET-based G protein dissociation assay. **(C)** Heatmap showing the inhibitory effects of the 48 candidate molecules on the force (3 pN)-induced Gs activation in LPHN2-overexpressing HEK293 cells measured by BRET-based G protein dissociation assay. Data are from 3 independent experiments (n = 3). **(D)** Dose-dependent inhibition of D11 on the force (3 pN)-induced Gs activation in LPHN2-overexpressing HEK293 cells (n = 3). **(E)** The chemical structure of the LPHN2 inhibitor D11. **(F-G)** The binding pose of D11 in the ligand pocket of LPHN2 (F) and the detailed interactions between D11 and LPHN2 pocket residues (G) according to computational simulation. **(H)** Effects of alanine mutation of LPHN2 pocket residues on the inhibitory potency of D11. The heatmap was generated according to the differences of inhibitory potency (ΔpEC50) of D11 between wild-type LPHN2 and its mutants. Data are from 3 independent experiments (n = 3). ND, no detectable inhibition. **(I)** Inhibitory effects of D11 on force (3 pN)-induced or ligand-induced G protein activation in HEK293 cells transfected with distinct combinations of aGPCR and G protein probes. Data are from 3 independent experiments (n = 3).

### LPHN2 regulates the MET current in a Gs-independent manner

We next exploited the reversible LPHN2-specific inhibitor D11 to investigate the immediate mechanism by which LPHN2 contributes to MET currents in OHCs from the mid-apical turn of WT cochlea in response to fluid jet stimulation. Since recent transcriptomic and functional studies at the single-cell level revealed heterogeneity among cochlear hair cells, which might be correlated with the tonotopic mapping of the cochlea (Frank et al., 2009; Petitpre et al., 2022; Shi et al., 2023), we also examined the expression levels of key components, including *Lphn2*, *Tmc1* and *Gnas*, in each of these tested cells by single-cell qRT-PCR (Figure 6A). Despite the fact that the baseline MET currents were normal, none of the *Lphn2*-negative hair cells (18.7% of the total) exhibited any detectable response to treatment with 50 nM D11, which was consistent with the high selectivity of D11 that was observed in the in vitro biochemical assays (Figures 6B and 6C). Intriguingly, among *Lphn2*-positive hair cells, hair cells that exclusively expressed *Lphn2* but not those that expressed *Tmc1* (*Lphn2+Tmc1-*, 5.3% of the total) showed no significant response to D11 treatment (Figures 6B and 6C). In contrast, MET currents in hair cells that expressed both *Lphn2* and *Tmc1* (*Lphn2+Tmc1+*, 76% of the total) were reduced by approximately 35% in response to treatment with 50 nM D11 and returned to normal after D11 was washed out (Figures 6B-6C and S6A-S6B). These results not only suggest that LPHN2 participates in the MET process but also suggest an interplay between LPHN2 and TMC1 in force sensation. Consistent with this hypothesis, the inhibitory effects of D11 on the MET response to fluid jet stimulation were abolished in cochlear hair cells that were harvested from *Pou4f3-CreER^+/-^Lphn2^fl/fl^* mice or from *Tmc1^-/-^* mice (Figures 6D and 6E).

**Figure 6.**
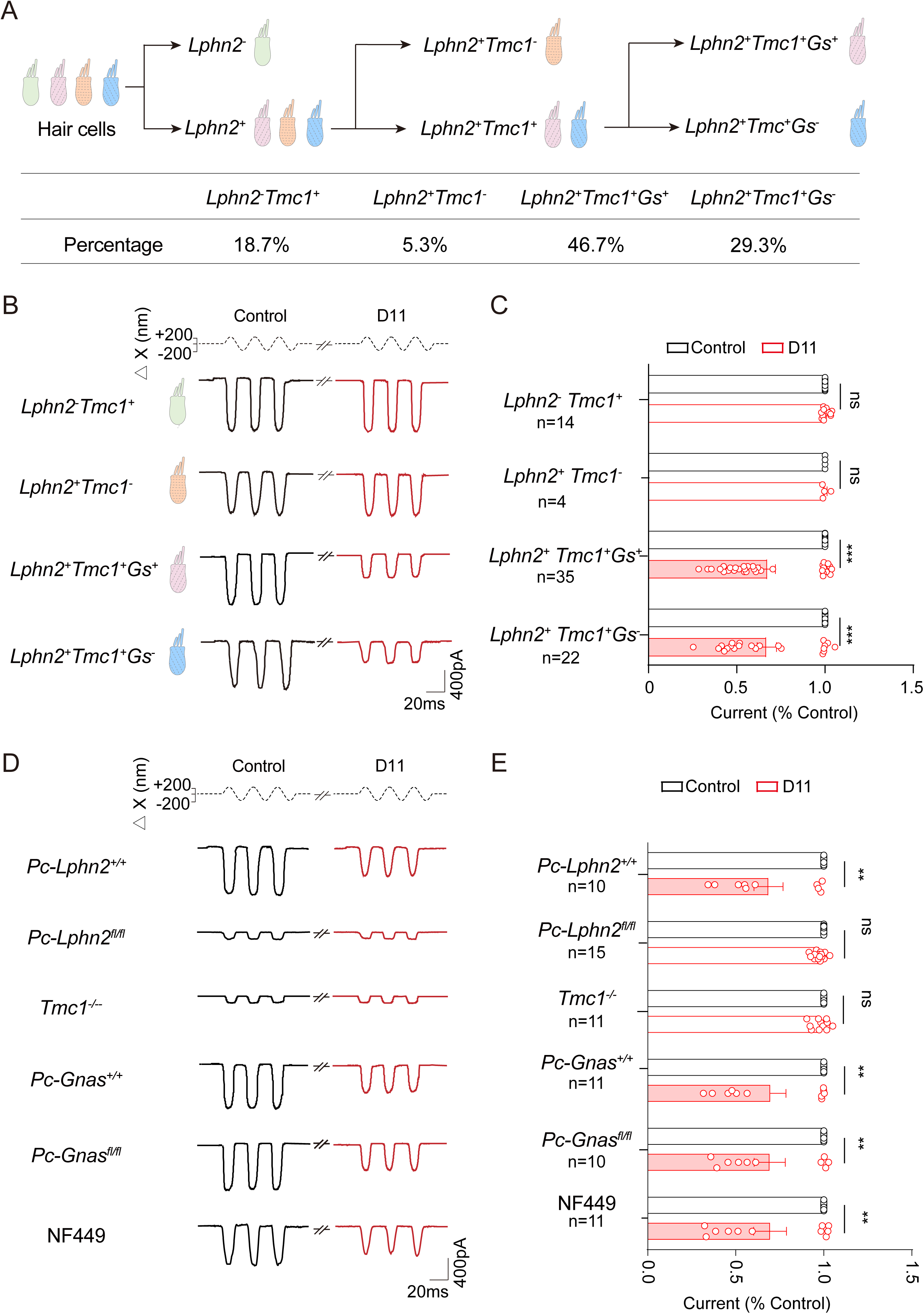
LPHN2 regulates MET currents in TMC1-dependent and Gs-independent manner. **(A)** schematic representation (upper panel) and quantitative analysis (bottom panel) of cochlear hair cell subclasses according to the expression of *Lphn2*, *Tmc1*, and *Gnas*. Data are correlated to Figure 6B-6C. **(B-C)** Representative current traces (B) and quantitative analysis (C) of fluid jet-stimulated MET responses of distinct cochlear hair cell subclasses pretreated with control vehicle or 50 nM D11. Data are normalized to the saturated MET response of control vehicle-treated hair cells in respective subclasses (n = 14, 4, 35, and 22 for *Lphn2^-^Tmc1^+^*, *Lphn2^+^Tmc1^-^*, *Lphn2^+^Tmc1^+^Gnas^+^* and *Lphn2^+^Tmc1^+^Gnas^-^* cells, respectively). Data are correlated to Figure S6C. **(D-E)** Representative current traces (D) and quantitative analysis (E) of fluid jet-stimulated MET responses of cochlear hair cells derived from *Pc-Lphn2^+/+^*, *Pc-Lphn2^fl/fl^*, *Pc-Gnas^+/+^*, or *Pc-Gnas^fl/fl^*mice in the absence or presence of 50 nM D11 or 20 μM NF449. Data are normalized to the saturated MET response of control vehicle-treated hair cells in respective groups (n = 10, 15, 11, 11, 10 and 11 for *Pc-Lphn2^+/+^*, *Pc-Lphn2^fl/fl^*, *Tmc1^-/-^*, *Pc-Gnas^+/+^*, *Pc-Gnas^fl/fl^* or NF449-treated *Pc-Gnas^+/+^* cells, respectively). Data are correlated to Figure S6F-S6G. **(C, E)** **P < 0.01; ***P < 0.001; ns, no significant difference. Cochlear hair cells pretreated with D11 compared with those treated with control vehicle. The bars indicate mean ± SEM values. All data were statistically analysed using paired two-sided Student’s *t* test.

We next explored the potential contribution of the Gs-cAMP pathway to the LPHN2-regulated MET response. Notably, no significant difference was observed between the baseline MET currents of Gs-positive (*Lphn2+Tmc1+Gnas-*, ∼30% of the total) and Gs-negative (*Lphn2+Tmc1+Gnas+*, ∼45% of the total) hair cells (718.7 pA vs. 716.9 pA) or between the D11-induced reduction in MET currents in these two types of cells (50% vs. 51%) (Figures S6C and S6D). Therefore, LPHN2 may regulate MET currents in a Gs-independent manner. Consistently, neither the genetic ablation of Gs in *Pou4f3-CreER^+/-^Gnas^fl/fl^* mice nor the pharmacological inhibition of Gs activity with the nonspecific Gs inhibitor NF449 significantly affected the inhibitory effects of D11 on the MET currents in cochlear hair cells (Figures 6D-6E and S6E-S6G). Collectively, these data suggest that the acute MET process that is regulated by the mechanosensitive receptor LPHN2 occurs in a TMC1-dependent and Gs-cAMP-independent manner.

### Colocalization and association of LPHN2 with the TMC1 complex

To further delineate the mechanism by which LPHN2 participates in the MET process, we next performed proteomic analyses to investigate the interactome of LPHN2 in mouse cochleae. The Flag-tagged LPHN2-GAIN construct showed mechanosensitive potential that was comparable to that of full-length LPHN2 (Figures 7A, S7A and S7B). We then purified the Flag-tagged LPHN2-GAIN protein from the baculovirus-*Spodoptera frugiperda* (Sf9) insect cell system and used the purified receptor as bait to search for potential candidates that coordinate LPHN2 function in vivo (Figure 7A). Notably, several components of the MET channel complex, including TMC1 and PCDH15, were detected in the complexes that were precipitated from cochlear homogenates by LPHN2-GAIN-bait but not in those that were precipitated by control bait (empty anti-Flag M2 beads) (Figures 7B and 7C). The interactions between LPHN2 and the proteins mentioned above as well as other known MET channel components, such as TMIE and LHFPL5, were further supported by an in vivo coimmunoprecipitation assay that was performed with commercially available antibodies (Figure 7D). Our in vitro coimmunoprecipitation experiments using a HEK293 cell overexpression system further confirmed the direct interaction between LPHN2 and TMC1 (Figure S7C). Moreover, LPHN2 exhibited significant coimmunostaining with these MET channel components at the shorter rows of stereocilia in cochlear hair cells at P10, suggesting that LPHN2 may directly associate with the MET channel complex *in vivo* (Figures 7E and 7F).

**Figure 7.**
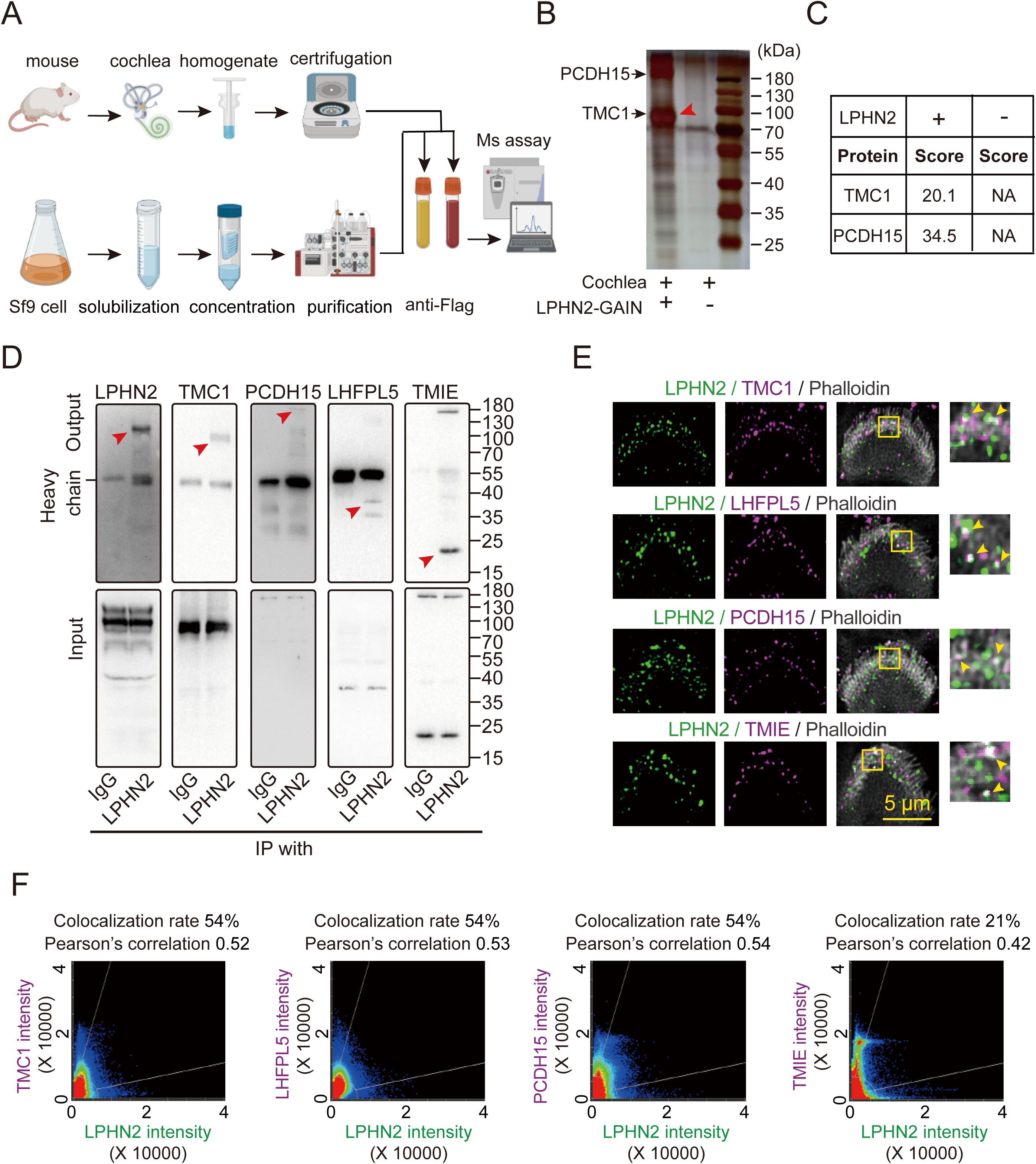
LPHN2 colocalizes and associates with MET channel components. **(A)** Schematic representation of the LPHN2-GAIN purification and mass spectrometry-based proteomic strategies used to identify potential LPHN2 interacting proteins in mouse cochlea. **(B)** Representative gel image showing the interacting proteins pulled-down by LPHN2-GAIN-batis or control baits (empty anti-Flag M2 beads). Red arrow indicates the band with similar molecular weight as TMC1. **(C)** MET channel components (TMC1 and PCDH15) specifically detected in cochleae homogenate pulled-down by LPHN2-GAIN-baits but not in that pulled-down by control baits. Data are correlated to Table S2. **(D)** Co-immunoprecipitation of LPHN2 with MET channel components (TMC1, PCDH15, LHFPL5 and TMIE) in the lysates of mouse cochlea. Representative blotting from three independent experiments is shown (n = 3). **(E)** Co-immunostaining of LPHN2 (green) with MET channel components (magenta) in the stereocilia of OHC in cochlear whole mounts derived from WT mice (n = 6 mice per group). Scale bar, 5 μm. Arrows indicate the colocalization of LPHN2 with MET channel components at the tips of shorter rows of stereocilia. **(F)** Pearson’s correlation analysis of the colocalization of LPHN2 with MET channel components. The Pearson’s correlation coefficient was 0.52, 0.53, 0.54 and 0.42 for the colocalization of LPHN2 with TMC1, LHFPL5, PCDH15 and TMIE, respectively. Data are correlated to Figure 7E.

### Functional coupling of the mechanosensitive receptor LPHN2 to TMC1

The potential association of LPHN2 with TMC1, which is the putative pore-forming subunit of the MET channel, prompted us to further investigate the functional interplay of these two membrane proteins. Thus, we attempted to reconstitute the functional coupling of LPHN2 and TMC1 in the heterologous HEK293 cell system. However, it is well known that TMC1 does not traffic properly to the plasma membrane when it is expressed in cultured cells (Beurg et al., 2015; Jia et al., 2020; Zhao et al., 2014). GPCRs are known to transport or recruit many cytoplasmic proteins, including but not limited to G proteins, arrestins, GRKs and especially PDZ domain-containing proteins, to the cell membrane of cochlear hair cells (Guan et al., 2023; Hall et al., 1998; Komolov et al., 2017; Shukla et al., 2014). We therefore screened the ability of the top 50 most highly expressed GPCRs in cochlear hair cells to transport TMC1 to the plasma membrane. Interestingly, two GPCRs, namely, GPCR-hc1 and GPCR-hc2, were found to transport TMC1 to the cell membrane when individually coexpressed with TMC1 in HEK293 cells, and GPCR-hc1 showed greater efficiency than GPCR-hc2 (Figures 8A, 8B and S7D). These two GPCRs that were identified provided useful tools for further investigating the electrophysiological properties of the interaction between LPHN2 and TMC1 in vitro.

**Figure 8.**
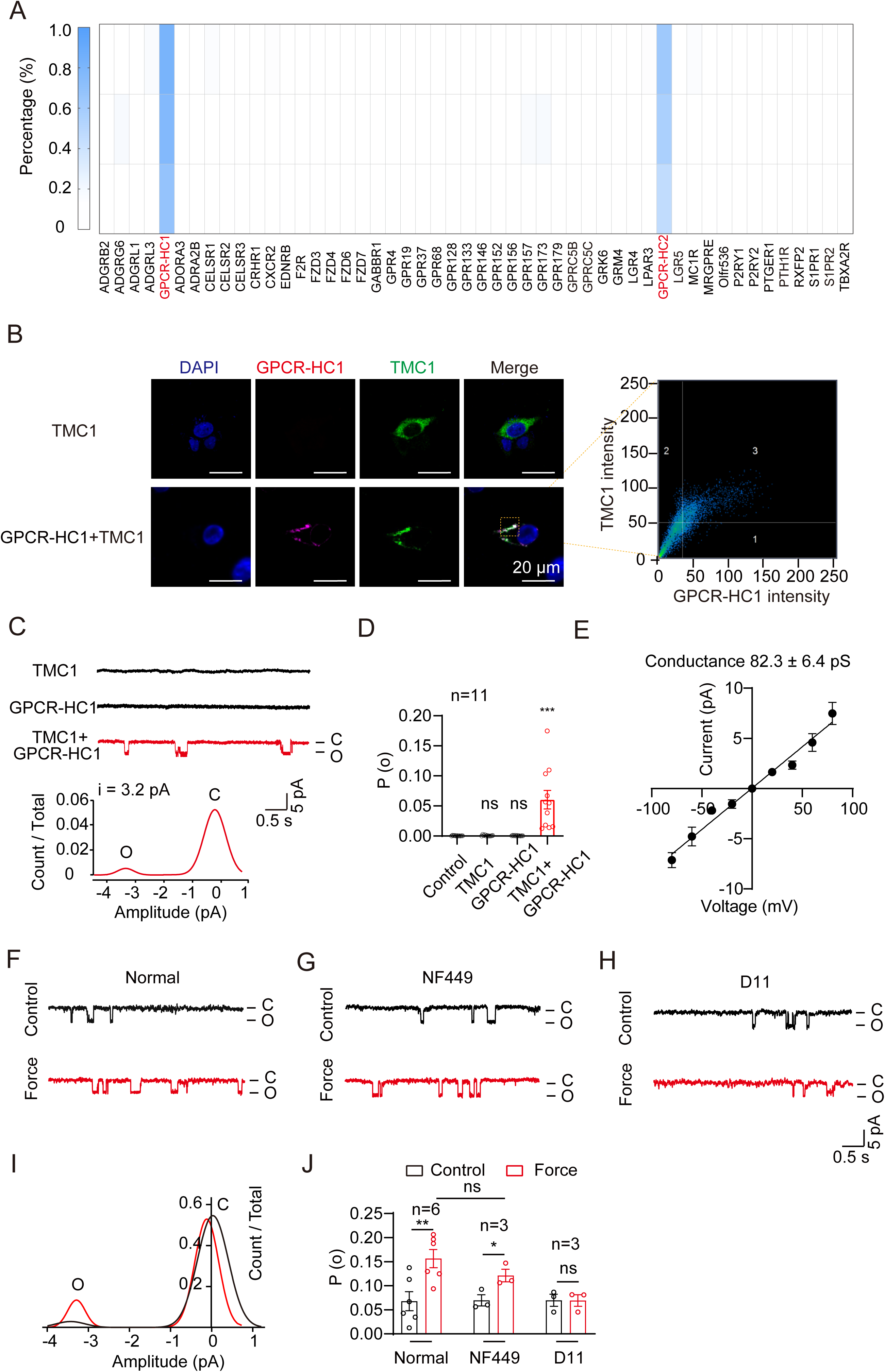
LPHN2 regulates MET through coupling to TMC1. **(A)** Heatmap showing the efficiency of top 50 abundantly-expressed GPCRs in cochlear hair cells for trafficking the TMC1 onto the plasma membrane. The efficiency was calculated as the ratio of TMC1 immunofluorescence intensity on plasma membrane to that in cytosol in HEK293 cells co-transfected with TMC1 and GPCR. Data are from 3 independent experiments (n = 3). **(B)** Left panel: Co-immunostaining of TMC1 (green) with GPCR-hc1 (magenta) in HEK293 cells transfected with TMC1 only or with TMC1 and GPCR-hc1. Scale bar: 20 μm. Representative images from three independent experiments were shown (n = 3). Right panel: Pearson’s correlation analysis of TMC1 and GPCR-hc1 fluorescence intensities. The Pearson’s correlation coefficient was 0.62. **(C)** Upper panel: Representative spontaneous single-channel currents of TMC1 at -40 mV recorded in HEK293 cells transfected with only TMC1 or GPCR-hc1, or with TMC1 and GPCR-hc1. Bottom panel: The normalized all-point amplitude histogram analysis of single-channel currents in HEK293 cells transfected with TMC1 and GPCR-hc1. The distribution data were fitted by a sum of two Gaussians, and the peaks correspond to the closed (C) and open (O) states. **(D)** Scatterplots of the single-channel open probability (Po) in HEK293 cells transfected with only TMC1 or GPCR-hc1, or with TMC1 and GPCR-hc1 (n = 11 per group). Data are correlated to Figure 8C. **(E)** The current-voltage relationship of the spontaneous currents recorded in HEK293 cells transfected with TMC1 and GPCR-hc1 (n = 3). The The single-channel conductance of TMC1 was determined as 82.3 ± 6.4 pS. Data are correlated to Figure S7E. **(F-H)** Representative spontaneous and force (10 pN)-stimulated single-channel current traces recorded in HEK293 cells transfected with TMC1 and GPCR-hc1 pretreated with control vehicle (F), 10 μM NF449 (G), or 50 nM D11(H). Data are correlated to Figure 8I-8J. **(I)** Histogram showing single-channel open probability at rest condition (black) and in response to 10 pN force stimulation (red). Data are correlated to Figure 8F. **(J)** Effect of NF449 or D11 treatment on the force-induced single-channel open probability (n = 6, 3 and 3 for control vehicle, NF449 and D11 treated group, respectively). Data are correlated to Figure 8F-8H. **(D)** ***P < 0.001; ns, no significant difference. HEK293 cells transfected with TMC1 or GPCR-hc1, or with TMC1 and GPCR-hc1 compared with the control cells. (**J**) *p < 0.05; **P < 0.01. Force-stimulated cells compared with control cells. ns, no significant difference. NF449-treated cells compared with control vehicle-treated cells. The bars indicate mean ± SEM values. All data were statistically analyzed using one-way (D) or two-way (J) ANOVA with Dunnett’s post hoc test.

We next performed patch-clamp recording on HEK293 cells that were cotransfected with GPCR-hc1 and TMC1, and we observed spontaneous single-channel opening at a holding potential of -40 mV (Figure 8C). As a negative control, HEK293 cells that were transfected with only TMC1 or GPCR-hc1 showed no detectable single-channel events, thus excluding the possibility that the recordings were from other unknown ion channels potentially mobilized by GPCR-hc1 (Figure 8C). The all-point histogram of single-channel opening events indicated that the average single-channel TMC1 current was 3.6 ± 0.2 pA, and the probability of TMC1 opening was 0.06 ± 0.01 (Figure 8D). The spontaneous single-channel events in HEK293 cells that coexpressed TMC1 and GPCR-hc1 were voltage dependent when a series of voltage steps from -100 mV to +100 mV were applied (Figures 8E and S7E). The average conductance was determined to be 82.3 ± 6.4 pS, which was similar to that reported for mouse TMC1 in apical cochlear hair cells in vivo (84.8∼88 pS) (Beurg et al., 2018; Fettiplace et al., 2022) (Figure 8E). Importantly, spontaneous single-channel events in HEK293 cells that coexpressed TMC1 and GPCR-hc1 were markedly inhibited or abrogated by pretreatment with known MET channel blockers, such as FM1-43 (3 μM), amiloride (0.2 mM) or dihydrostreptomycin (0.2 mM) (Figure S7F). These experimental results collectively suggested that TMC1 was rendered functional by coexpression with GPCR-hc1 in the heterologous HEK293 cell system.

To further examine whether the sensing of force by LPHN2 regulates the activity of TMC1, we applied LPHN2-M-beads to the heterologous HEK293 cell reconstitution system coexpressing TMC1, LPHN2 and GPCR-hc1, and we recorded single-channel opening events in response to the application of 10 pN force in a magnetic field; this force equals to 100 dynes/cm^2^ on the plasma membrane and mimics physiological stimulation at a velocity of approximately 1 m/s. By resolving the single-channel currents, we observed an approximately 2.2-fold increase in the probability of TMC1 opening in HEK293 cells that coexpressed GPCR-hc1, LPHN2 and TMC1 in response to force stimulation compared with that resting conditions (Figures 8F and 8J). However, the single-channel current amplitude was not significantly affected by the administration of force via LPHN2-M-beads (Figure S7G). As a negative control, no force-induced single-channel events were observed in HEK293 cells that coexpressed GPCR-hc1 with LPHN2 or TMC1 alone or in cells that coexpressed GPCR-hc1 with both CDH23 and TMC1 (Figures S7H and S7I). Importantly, the LPHN2 mediated force-induced single-channel opening was abolished by pretreatment with the LPHN2-specific inhibitor D11 (50 nM) or the TMC1 inhibitor amiloride (0.2 mM), but it was not significantly affected by pretreatment with the nonspecific Gs inhibitor NF449 (20 μM) (Figures 8G-8J and S7J-S7K). Intriguingly, an approximately 1.6-fold increase of single-channel currents was also observed in HEK293 cells co-expressing GPCR-hc1, PCDH15 and TMC1 in response to 10 pN force stimulation via PCDH15-M-beads, but not in samples stimulated with control beads (Figures S7M and S7N). The TMC1 current was increased by approximately 2.6-fold when LPHN2 was further co-expressed in this system (GPCR-hc1, PCDH15 and TMC1) (Figures S7M and S7N). Therefore, these in vitro data showing that LPHN2 activates TMC1 in response to force stimulation are consistent with the functional and colocalization data from the in vivo experiments, suggesting that LPHN2 can convert extracellular mechanical stimuli to activate TMC1 in a Gs-independent manner. The LPHN2-mediated moderate increase of TMC1 open probability and its up-regulatory effect on PCDH15-induced TMC1 activation collectively imply that LPHN2 may be a component of MET machinery that modulate the tip-link-mediated MET process in sound sensation.

### Force sensation by LPHN2 induces conformational changes in TMC1

The results of colocalization and co-IP of LPHN2 and TMC1 suggested that LPHN2 directly interacts and forms a complex with TMC1. To elucidate the mode of interaction between the seven transmembrane receptor LPHN2 and the ten transmembrane ion channel TMC1, we performed a BRET assay by inserting Nanoluc luciferase (Nluc) at specific positions in the intracellular loops (ICLs) or C-terminus of LPHN2 (denoted as LPHN2-ICL1/2/3-Nluc or LPHN2-C-Nluc, respectively) and adding a YFP tag to the N- or C-terminus of TMC1 (denoted as TMC1-N-YFP and TMC1-C-YFP, respectively) (Figure 9A). This method has been used to characterize the interaction between GPCRs or between GPCRs and regulatory molecules in previous studies (Angers et al., 2000; Fu et al., 2021; Guan et al., 2023; Harikumar et al., 2012). Specific saturation BRET signals were detected between LPHN2-ICL1/2-Nluc and TMC1-C-YFP in HEK293 cells (coexpressing GPCR-hc1 as described above), and these signals depended on the channel expression level; however, such signals were not detected between the other BRET pairs, suggesting that the ICL1/2 (or potentially TM1-4) region of LPHN2 is in proximity to the C-terminus or TM10 region of TMC1 (Figures 9B-9C and S8A). Moreover, the application of force to LPHN2 increased the BRET signal between LPHN2-ICL1-Nluc and TMC1-C-YFP in a dose-dependent manner, with an EC50 of 5.2 ± 1.1 pN; this value was similar to the EC50 of force-induced TMC1 activation via LPHN2 in the heterologous system (Figures 9D and 9E). Notably, the BRET_50_ value for LPHN2-Nluc and TMC1-YFP was recorded when the interacting pairs were expressed at a 1:2 expression ratio, suggesting a 1:2 stoichiometry in the LPHN2-TMC1 heteromer.

**Figure 9.**
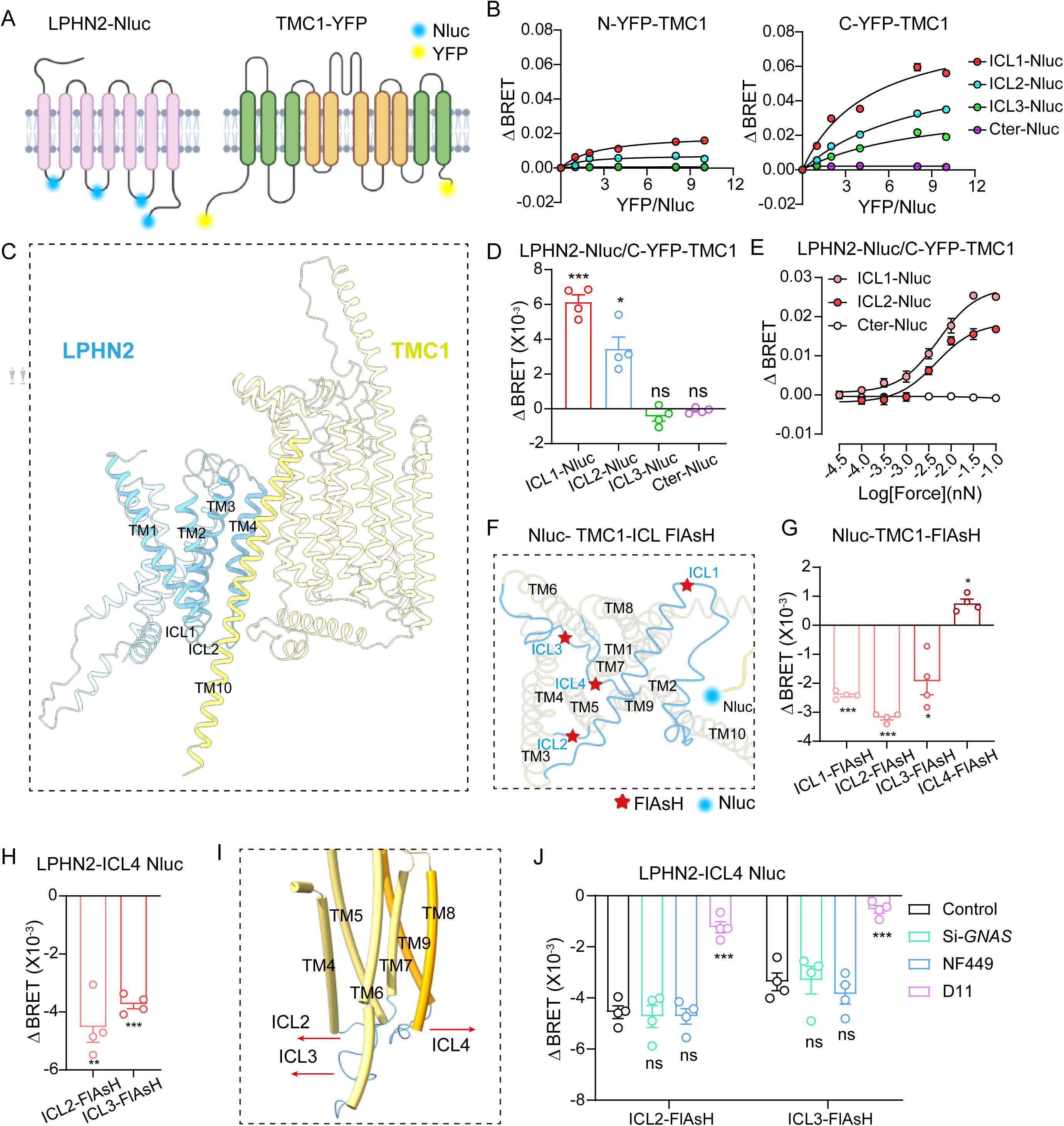
LPHN2 converts force stimuli into the conformational change of TMC1. **(A)** Schematic diagram showing the generation of BRET pairs by fusing YFP at the N- or C-terminus of TMC1 (referred to as N-YFP-TMC1 or C-YFP-TMC1) and inserting Nluc into specific positions of different ICLs of LPHN2 (referred to as L2-ICL1-Nluc, L2-ICL2-Nluc, L2-ICL3-Nluc and L2-Cter-Nluc, respectively). **(B)** Saturation BRET signals between NLuc and YFP in HEK293 cells co-transfected with a fixed amount of NLuc-tagged LPHN2 and an increasing amount of TMC1-YFP. Data are from three independent experiments (n = 3). Data are correlated to Figure S8A. **(C)** Interaction model of LPHN2 and TMC1 according to computational simulation. **(D)** Force-induced intermolecular BRET signals between C-YFP-TMC1 and Nluc-tagged LPHN2 (n = 4). **(E)** Force-induced dose-dependent BRET signal increase between C-YFP-TMC1 and Nluc-tagged LPHN2 (n = 3). **(F)** Schematic diagram showing the generation of TMC1 conformational sensors by fusing the Nluc at the N-terminus of TMC1 and inserting the FlAsH probes (CCPGCC) into specific positions of different ICLs of TMC1. **(G)** Intramolecular BRET signals between the Nluc and FlAsH probes in the respective Nluc-TMC1-FlAsH constructs in response to the force stimulation on LPHN2 (n = 4). Data are correlated to Figure S8B. **(H)** Intramolecular BRET signals between the Nluc fused in ICL4 of TMC1 and FlAsH probes inserted at ICL2 or ICL3 in response to the force stimulation on LPHN2 (n = 4). Data are correlated to Figure S8C. **(I)** Schematic diagram showing the movement of ICL2/3 and ICL4 or TMC1 towards the opposite direction in response to force stimulation on LPHN2. **(J)** Effects of pretreatment with si*GNAS*, NF449 (10 μM) or D11 (50 nM) on the force-induced intramolecular BRET signals between the Nluc fused in ICL4 of TMC1 and FlAsH probes inserted at ICL2 or ICL3 (n = 4). (**D, G, H**) *P < 0.05; **P < 0.01; ***P < 0.001; ns, no significant difference. Force-stimulated HEK293 cells compared with the control cells. **(J)** ***P < 0.001; ns, no significant difference. HEK293 cells treated with si*GNAS*, NF449 or D11 compared with those treated with control siRNA or control vehicle. The bars indicate mean ± SEM values. All data were statistically analyzed using unpaired two-sided Student’s *t* test (D, G, H) or one-way ANOVA with Dunnett’s post hoc test (J).

To investigate force-induced conformational changes in TMC1, we added Nluc to the N-terminus of TMC1 and inserted FlAsH probes (CCPGCC) into specific positions of the ICLs to generate a panel of TMC1 conformational sensors (Figure 9F). Such conformational sensors are often used to probe conformal changes in membrane proteins (Fu et al., 2021; Lin et al., 2022; Yang et al., 2021; Yang et al., 2024). In response to the application of force on LPHN2, the intramolecular BRET signals between the N-terminus and the ICL1, ICL2, or ICL3 of TMC1 decreased, suggesting the dissociation of these ICLs from the N-terminus (Figures 9G and S8B). In contrast, a significant increase in the BRET signal was observed at ICL4, indicating that this specific structural element moved close to the N-terminal Nluc (Figure 9G). A similar result was also observed when Nluc was fused to the ICL4 of TMC1 and the FlAsH probe was inserted in the ICL2 or ICL3; these data suggested that a separation between ICL2/3 and ICL4 of TMC1 in response to the application of force to LPHN2 (Figures 9H and S8C). Importantly, the cryo-EM structure of the *C. elegans* TMC1 complex suggests that the putative ion-conduction pathway of TMC1 mainly includes TM4-TM8 (Jeong et al., 2022). Here, we observed that in TMC1, ICL2/3 moved in an opposite direction compared with ICL4 in response to force sensation by LPHN2. This conformational change in TMC1 indicates a separation between the TM4-7 and TM8-9 ends, which indicates the potential opening of the ion channel pore (Figure 9I). RNAi-mediated knockdown of Gs or administration of the nonspecific Gs inhibitor NF449 had no significant effect on the conformational changes in TMC1. In contrast, the LPHN2-specific inhibitor D11 suppressed these conformational changes (Figure 9I). Collectively, our results suggest that LPHN2 converted force stimulation into conformational changes in TMC1 in a Gs-independent manner, and this effect was potentially mediated through physical interactions that led to the opening of the ion-conduction pore of TMC1.

### Force sensation by LPHN2 stimulates a Ca^2+^ response and neurotransmitter release in cochlear hair cells

It is suggested that in response to acoustic stimulation, neurotransmitters such as glutamate are released from the ribbon synapse and regulate the excitability of spiral ganglion neurons (SGNs), which transmit sound information from cochlear hair cells to the central nervous system (Qiu and Muller, 2022; Sun et al., 2018). To explore the potential effects of force stimulation via LPHN2 on glutamate release from cochlear hair cells, we employed the red fluorescent glutamate sensor R^ncp^-iGluSnFR (R^ncp^-iGlu) under the control of the *hSyn* promoter, which enables specific expression of this sensor in SGNs and recording of glutamate release in real time (Figure 10A) (Wu et al., 2018). Notably, repeated application of force to the cochlear sensory epithelium by LPHN2-M-beads resulted in marked glutamate secretion, with an EC50 of 5.1 ± 0.9 pN, and this glutamate secretion was inhibited by pretreatment with the LPHN2-specific inhibitor D11 (Figures 10B-10D). As a negative control, the force that was applied by Ctrl-beads to WT hair cells or by LPHN2-M-beads to hair cells from *Pou4f3-CreER^+/-^Lphn2^fl/fl^*mice did not induce any detectable glutamate secretion (Figures S9A and S9B).

**Figure 10.**
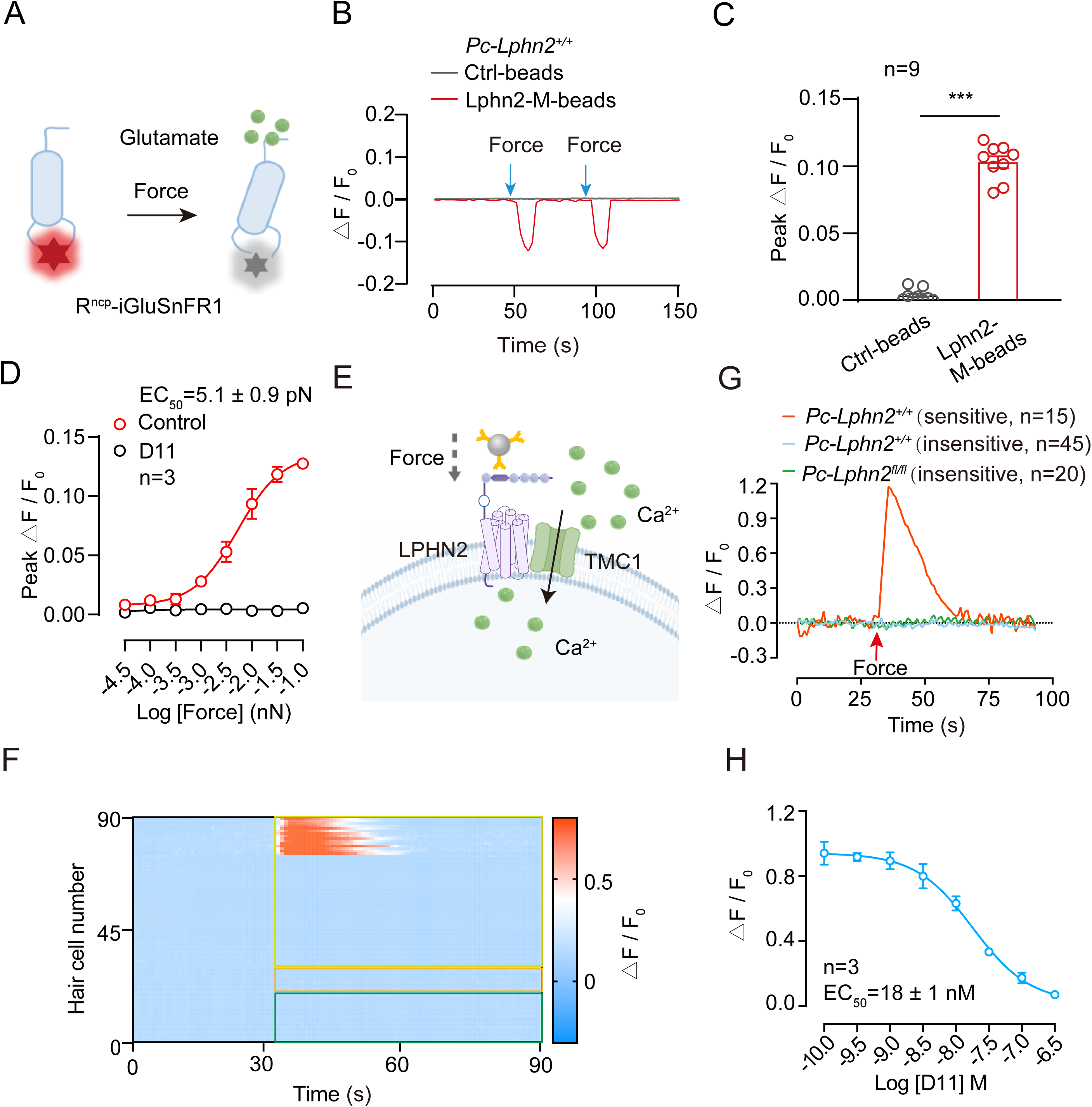
Force sensation by LPHN2 induces glutamate release and Ca^2+^ signals in cochlear hair cells. **(A)** Schematic diagram showing the working mechanism of the red fluorescent glutamate sensor Rncp-iGluSnFR (Rncp-iGlu). **(B-C)** Representative traces (B) and quantitative analysis (C) of glutamate secretion from individual cochlear hair cell derived from the P10 *Pc-Lphn2^+/+^* mice in response to force applied by LPHN2-M-beads or control beads (n = 9). Data are correlated to Figure S9A-S9B. **(D)** Force-induced dose-dependent glutamate secretion from cochlear hair cells in the absence or presence of D11. Representative curves from three independent experiments are shown (n = 3). **(E)** Schematic illustration of the Ca^2+^ imaging in mouse cochlea hair cells labelled by AAV-ie-*L2pr*-mCherry in response to force stimulation. **(F)** Heatmaps showing the Ca^2+^ responses in individual cochlear hair cell derived from the P10 *Pc-Lphn2^+/+^* or *Pc-Lphn2^fl/fl^* mice. n=60 for the AAV-ie-*L2pr*-mCherry-labelled cochlear hair cells from *Pc-Lphn2^+/+^* mice (yellow frame); n=10 for the unlabelled hair cells from *Pc-Lphn2^+/+^* mice (orange frame); n = 20 for the AAV-ie-*L2pr*-mCherry-labelled cochlear hair cells from *Pc-Lphn2^fl/fl^* mice (green frame). Y axis represents the magnitude of the calcium response characterized by △F/F_0_. **(G)** Representative Ca^2+^ traces recorded in individual cochlear hair cells derived from P10 *Pc-Lphn2^+/+^* or *Pc-Lphn2^fl/fl^*mice. **(H)** Dose-dependent inhibitory effects of D11 treatment on force-stimulated Ca^2+^ responses in cochlear hair cells labelled by AAV-ie-*L2pr*-mCherry. Data are correlated to Figure S9C. **(C)** ***P < 0.001. Cochlea explants treated with LPHN2-M-beads compared with those treated with Ctrl-beads. The bars indicate mean ± SEM values. Data were statistically analyzed using unpaired two-sided Student’s *t* test.

Neurotransmitter release from cochlear hair cells is often regulated by intracellular Ca^2+^ concentrations; these concentrations are hypothesized to increase locally near the afferent synapse in response to mechanical stimulation via the formation of Ca^2+^ nanodomains (Jaime Tobon and Moser, 2023; Meyer et al., 2009). To record the force-induced Ca^2+^ response at the single-cell level, P10 cochleae from mice that were treated with modified AAV-ie-*Lphn2pr*-mCherry were isolated, and LPHN2-expressing (mCherry-labeled) hair cells were selected for Ca^2+^ imaging using a Fura-2 probe (Figure 10E). Approximately 25% of the LPHN2-expressing OHCs exhibited a significant Ca^2+^ response to LPHN2-M-bead-mediated force stimulation (the response rate might have been affected by the efficiency with which the LPHN2-M-beads bound to endogenous LPHN2 on stereocilia) (Figures 10F and 10G). In contrast, neither the unlabeled cochlear hair cells nor the mCherry-labeled hair cells derived from the *Pou4f3-CreER^+/-^Lphn2^fl/fl^*mice produced detectable Ca^2+^ signals in response to force application, suggesting the specificity of the mechanical force-mediated Ca^2+^ response via LPHN2 activation (Figures 10F and 10G). The force-stimulated Ca^2+^ response in LPHN2-expressing cochlear hair cells was inhibited by pretreatment with the LPHN2-specific inhibitor D11 in a concentration-dependent manner, and the EC50 value was 18 ± 1 nM; this result was similar to the inhibitory effect of D11 on force-stimulated LPHN2 activation in the heterologous system, suggesting the specificity of LPHN2 in mediating the force-induced Ca^2+^ response (Figures 10H and S9C-S9D). Notably, in our previous proteomic analyses, we found that in addition to its association with MET channel components, LPHN2 also interacted with other cation channels, such as CNGA3, whose activity is regulated by cyclic nucleotides. Therefore, upon the sensing of force, LPHN2 might also activate CNGA3 through Gs-dependent cAMP upregulation. However, further studies are warranted to evaluate the potential contribution of TMC1-dependent and/or Gs-dependent signaling to the force-induced Ca^2+^ response in cochlear hair cells through LPHN2 activation. Nevertheless, our results indicated that LPHN2 directly participates in the MET process in cochlear hair cells and converts force stimulation into an intracellular Ca^2+^ response, leading to neurotransmitter release and potential regulation of SNG excitability.

### LPHN2 re-expression in cochlear hair cells of *Lphn2*-deficient mice reverses hearing loss

To further validate the functional roles of LPHN2 in the peripheral auditory system, especially in cochlear hair cells, we next performed an in vivo rescue experiment. In particular, we delivered LPHN2-encoding AAV-ie into the cochleae of *Pou4f3-CreER^+/-^Lphn2^fl/fl^* mice and examined its effects on the auditory functions of these *Lphn2*-deficient mice. The LPHN2-GAIN construct, which lacked the 523 N-terminal residues but retaining force sensation ability (Figures S7A and 7B), was packaged into the AAV-ie-*Lphn2pr*-mCherry vector (referred to as AAV-ie-*Lphn2pr*-LPHN2) and delivered into P3 *Pou4f3-CreER^+/-^Lphn2^fl/fl^* mice by round window membrane injection (Figure S10A). LPHN2-GAIN was specifically expressed in the cochleae and vestibules, but not in the brain, of virus-treated *Pou4f3-CreER^+/-^Lphn2^fl/fl^* mice, as shown via western blotting (Figure S10B). In particular, LPHN2 was re-expressed at the tips of shorter rows of stereocilia in both cochlear OHCs and IHCs of AAV-ie-*Lphn2pr*-LPHN2-treated *Pou4f3-CreER^+/-^Lphn2^fl/fl^* mice (Figures S10C-S10E). In contrast, no LPHN2-GAIN expression was observed in the cochleae of *Pou4f3-CreER^+/-^Lphn2^fl/fl^* mice that were treated with the AAV-ie-*Lphn2pr*-mCherry control virus (Figures S10C-S10E).

We next assessed the auditory functions of mice that were subjected to LPHN2-GAIN or control gene delivery. Notably, *Lphn2*-deficient mice that were treated with AAV-ie-*Lphn2pr*-LPHN2 exhibited significantly improved performance in the ABR test. The ABR thresholds at all the testing frequencies, as well as the amplitude and latency of ABR wave I, of AAV-ie-*Lphn2pr*-LPHN2-treated *Pou4f3-CreER^+/-^Lphn2^fl/fl^*mice were restored to levels that were comparable to those of *Pou4f3-CreER^+/-^Lphn2^+/+^*mice (Figures 11A-11C). Moreover, the DPOAE thresholds of the *Lphn2*-deficient mice were also restored to nearly normal levels after AAV-ie-*Lphn2pr*-LPHN2 treatment (Figure 11D). In contrast, *Lphn2*-deficient mice that were treated with AAV-ie-*Lphn2pr*-mCherry showed no significant improvements in any of the parameters mentioned above (Figures 11A-11D). The MET characteristics of LPHN2-expressing cochlear hair cells in response to fluid jet stimulation were also evaluated. Our results indicated that mCherry-labeled cochlear hair cells in AAV-ie-*Lphn2pr*-LPHN2-treated *Pou4f3-CreER^+/-^Lphn2^fl/fl^*mice showed MET currents with similar amplitudes and responsiveness to D11 as those that were recorded for LPHN2-expressing hair cells in WT cochleae (Figure 11E). In contrast, the severe impairments in the MET currents of LPHN2-expressing cochlear hair cells from *Pou4f3-CreER^+/-^Lphn2^fl/fl^* mice were not significantly ameliorated by treatment with the AAV-ie-*Lphn2pr*-mCherry control virus (Figure 11E). Collectively, these findings suggest that LPHN2 in cochlear hair cells directly contributes to auditory perception through the sensation of mechanical force.

**Figure 11.**
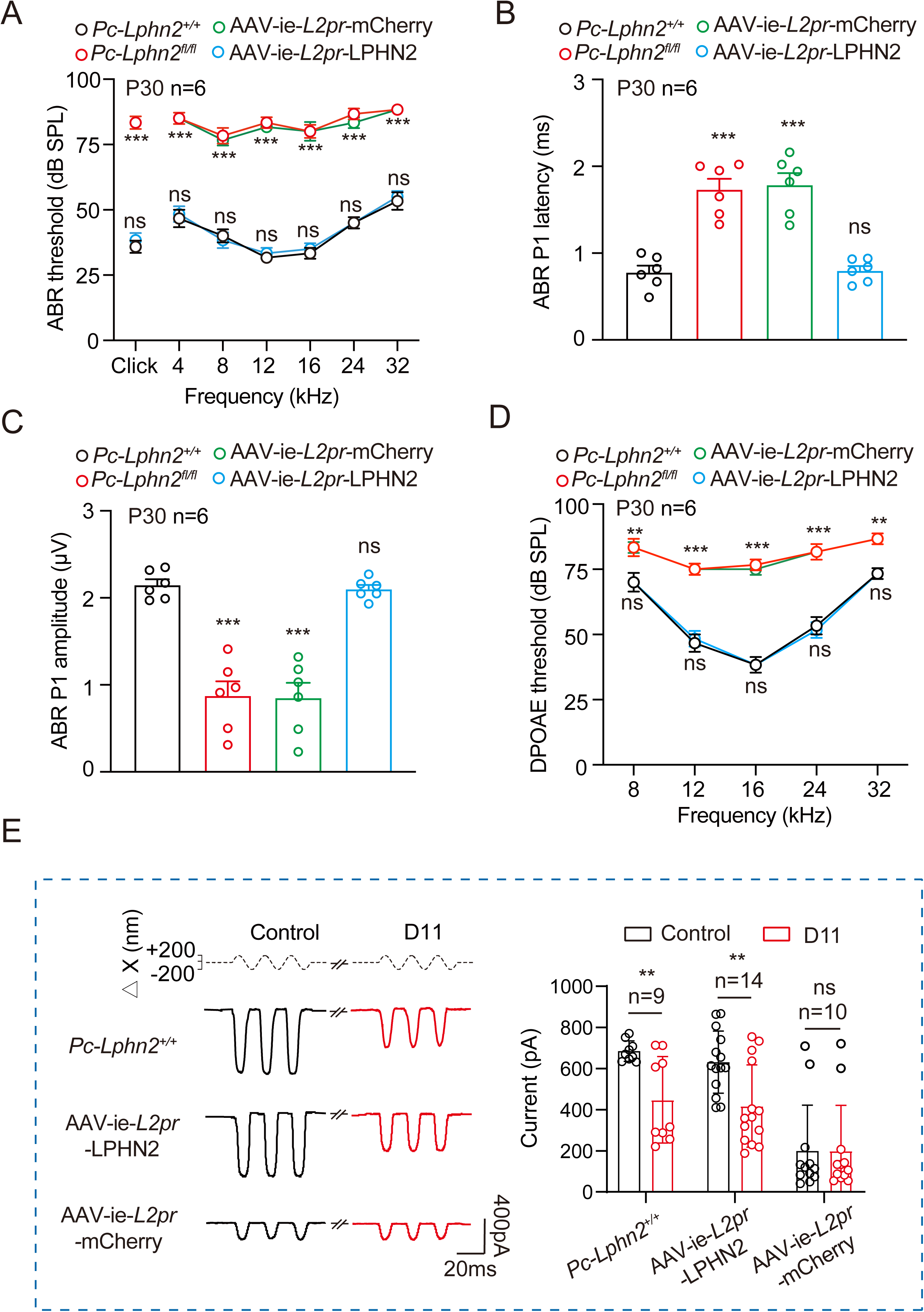
LPHN2 re-expression in cochlear hair cells of Lphn2-deficient mice rescues hearing loss and MET. **(A-D)** ABR thresholds (A), ABR peak1 (P1) latency (B) and amplitude (C), and DPOAE thresholds (D) of *Pc-Lphn2^+/+^*, *Pc-Lphn2^fl/fl^*, AAV-ie-*L2pr*-LPHN2-treated *Pc-Lphn2^fl/fl^* mice (referred to as AAV-ie-*L2pr-*LPHN2 mice) and AAV-ie-*L2pr*-mCherry-treated *Pc-Lphn2^fl/fl^*mice (referred to as AAV-ie-*L2pr-*mCherry mice) at P30 (n = 6 mice per group). Data are correlated to Figure S7A-S7E. **(E)** Representative current traces (left) and quantitative analysis (right) of fluid jet-stimulated MET responses of cochlear hair cells derived from *Pc-Lphn2^+/+^*, AAV-ie-*L2pr-*LPHN2 or AAV-ie-*L2pr-*mCherrymice in the absence or presence of 50 nM D11 (n = 9, 14 and 10 for *Pc-Lphn2^+/+^*, AAV-ie-*L2pr-*LPHN2 or AAV-ie-*L2pr-*mCherry cells, respectively). **(A-D)** **P < 0.01; ***P < 0.001; ns, no significant difference. *Pc-Lphn2^fl/fl^*, AAV-ie-*L2pr-* LPHN2 or AAV-ie-*L2pr-*mCherry mice compared with *Pc-Lphn2^+/+^* mice. **(E)** **P < 0.01; ns, no significant difference. Cochlea explants treated with D11 compared with those treated with control vehicle. The bars indicate mean ± SEM values. All data were statistically analyzed using one-way ANOVA with Dunnett’s post hoc test (A-D) or paired two-sided Student’s *t* test (E).

## Discussion

Cochlear hair cells convert sound stimuli into electrochemical signals through the activation of MET channels. A series of candidate components of the MET channel complex have been identified, among which TMC1/2 were proposed to be pore-forming subunits (Jia et al., 2020; Pan et al., 2018; Qiu and Muller, 2022). Because of the rapid activation and ultralow response latency of cochlear hair cells in the submillisecond range, the MET process is conventionally believed to be directly regulated by ion channels but not by GPCRs, which usually transduce signals through second messengers or enzymatic pathways (Corey and Hudspeth, 1979; Hudspeth, 2014). In the present work, we showed that a mechanosensitive GPCR, LPHN2, actively participates in the MET process in cochlear hair cells. Notably, LPHN2 is distributed at the tips of shorter rows of stereocilia in cochlear hair cells, where it colocalizes with MET channel components. In contrast to the signaling mechanism for smell or visual perception, which mainly relies on the secondary messenger system downstream of sensory GPCRs, LPHN2 directly interacts with TMC1 and plays an important modulatory role in the MET process during auditory perception; this conclusion was supported by genetic studies using *Lphn2*-deficient mice, pharmacological studies using a specific LPHN2 inhibitor and single-channel analyses in a heterologous reconstitution system. Importantly, our data suggested that the sensation of force by LPHN2 was converted to conformational changes in TMC1, increasing the probability of TMC1 opening. These data provide experimental evidence that a mechanosensitive GPCR might be a component of the MET complex. Interestingly, the *Drosophila* LPHN homolog, Ca^2+^-independent receptor of latrotoxin (drCirl), has been reported to respond to mechanical stimuli in vivo by interacting with the Toll-like receptor Tollo, which modulates the sensitivity of *Drosophila* to tactile and auditory stimuli (Scholz et al., 2023; Scholz et al., 2015). However, in contrast to the mechanism underlying the function of LPHN2 in mouse cochlear hair cells, Cirl converts mechanical stimulation to decrease the intracellular cAMP levels in the sensory neurons of *Drosophila* (Scholz et al., 2017). Whether drCirl directly couples to the *Drosophila* TMC homolog is worthy of future study. In addition, our parallel study indicated that another LPHN subfamily members, namely LPHN3, can also be activated in response to force stimulation, but heterozygous deficiency of this receptor resulted in no obvious balance or hearing disorder phenotype (unpublished observation). Therefore, we hypothesized that mechanosensitive LPHNs and their homologs might represent one subfamily of evolutionally conserved auditory sensors. Whether mechanistic differences contributed to faster and more accurate responses through evolutionary adaptation to sound stimuli in more complex environments requires further investigation.

Frequency, amplitude and period are the three most important variables of sound. Other variables may include timbre and envelope. In addition to hearing responses with very quick latency that are mediated by MET (required for frequency detection), signals ranging from 20 to 500 ms may also contribute to hearing experiences (may contribute to period detection). For example, our language communications do not always require very fast sensation/differentiation. People normally differentiate speech with similar voice frequencies at a rate of up to 9 syllables/second, indicating an approximately 110 ms differentiation time (Coupe et al., 2019; Peelle and Davis, 2012). Moreover, the calcium influx that occurs in hair cells in response to bundle deflection, which is a classic method for mimicking sound sensation ex vivo, can last approximately 200 ms (Beurg et al., 2009). Hearing responses with a latency of approximately 100 ms may play a regulatory role in responses to bundle deflection. These findings raised the interesting possibility that not only force sensors with a response latency in the microsecond range but also force sensors with a response latency ranging from 20 ms to 500 ms, as exemplified by GPCR-secondary messenger systems, could function as auditory modulators that regulate neurotransmitter release and hearing sensations. Here, in addition to its direct association with TMC1, our data suggested that LPHN2 activated Gs signaling and increased intracellular cAMP concentrations in cochlear hair cells in response to force stimulation. Previous studies suggested that cAMP signaling regulates the mechanotransduction process in rat cochlear hair cells and affects auditory sensitivity by regulating gating spring stiffness (Caprara and Peng, 2022; Mecca et al., 2022). Moreover, a loss-of-function mutation in adenylyl cyclase 1 (AC1), which is the upstream regulator of cAMP production, caused by the deletion of the 82 C-terminal amino acids has been associated with hearing loss in humans (Santos-Cortez et al., 2014). Therefore, Gs-cAMP signaling might also participate in auditory perception. However, our present work revealed that genetic ablation or pharmacological inhibition of Gs had no significant effect on the baseline MET of LPHN2-expressing hair cells or on their response to an LPHN2-specific inhibitor; these results suggest the dispensable role of Gs-cAMP signaling in the LPHN2-mediated regulation of MET. This result might be explained by the longer latency time of cAMP production (millisecond range) than that of acute MET (microsecond range). Nevertheless, we cannot exclude the possibility that force-induced Gs-cAMP signaling downstream of LPHN2 might also affect auditory perception by fine-tuning the sensitivity or adaptation of the MET process or by participating in longer-term regulation of hair cell physiology, such as neurotransmitter release or intercellular communication, via the regulation of cAMP-gated cation channel activity, such as CNG or HCN activity. Indeed, we have revealed an association between LPHN2 and CNGA3 in cochlear hair cells. Future studies using hair cell-specific Gs- or CNGA3-knockout mice combined with detailed activation/adaptation analyses of MET currents and neurotransmitter probing will be helpful for elucidating the functional roles of LPHN2-Gs-cAMP signaling in auditory perception.

We performed experiments to analyze the MET responses of cochlear hair cells expressing LPHN2 and that used the LPHN2 inhibitor D11, and our data suggested that auditory hair cells are heterogeneous in terms of force-sensitive GPCRs, such as LPHN2. Lphn2 was expressed in 80% of cochlear hair cells, and Lphn2 deficiency, specifically in *Pou4f3-CreER^+/-^Lphn2^fl/fl^*mice, caused hearing loss but not complete hearing loss. These results suggested that both Lphn2-positive and Lphn2-negative hair cells play important roles in hearing processes but that these hair cells are not functionally redundant. Specifically, although Lphn2-deficient hair cells exhibited severely impaired MET responses compared with those of control cells, LPHN2-negative hair cells in WT cochleae generally exhibited MET currents that were comparable to those of LPHN-positive hair cells (Figure S6C). Therefore, it is possible that other mechanosensitive components might exist in LPHN2-negative hair cells to compensate for the function of LPHN2 and regulate the MET response. There are many kinds of force, such as tension force, normal force, and friction force, which further include static friction force, sliding friction force, rolling friction force and fluid friction force. We indeed identified several force-sensing GPCRs other than LPHN2, which exhibit different expression patterns from those of LPHN2 in cochlear hair cells and also participate in the MET process (unpublished observation). At least one of these mechanosensitive GPCRs (msGPCR2) expressed on stereocilia was able to couple to TMC1. Similar to LPHN2, knockout or pharmacological inhibition of this msGPCR2 decreased MET in fluid jet assay. These observations may account for the less severe hearing loss phenotype observed in LPHN2-deficient mice compared with those with deficiency for other MET components, such as CIB2 or TMIE (data will be presented in another work)(Giese et al., 2017; Zhao et al., 2014). Currently, we do not know why hair cells need multiple GPCRs to sense mechanical stimuli. We hypothesized that these GPCRs might be sensitive to different mechanical stimuli in the cochlea or that these GPCRs might recognize different variables of sound, such as frequency, amplitude and period. However, further studies are needed to elucidate the detailed functions of these mechanosensitive GPCRs and to explore the potential effects of their combinations on auditory perception. Moreover, from a single-cell perspective, even the Lphn2^+^Tmc1^+^Gs^+^ cells exhibited heterogeneous MET responses to D11 treatment (approximately 30% of these cells were insensitive to D11), suggesting additional layers of complexity in hair cell heterogeneity (Figure 6C). This is potentially consistent with previous observations that SGNs driven by different cochlear hair cells could have different firing rates (Sun et al., 2018).

Considering the four criteria that we have proposed to fulfil the auditory receptor candidates, we found that Lphn2 is expressed in the stereocilia of cochlear hair cells (1); Lphn2 senses force with an EC50 of 1.7 ± 0.3 pN, which is within a physiological range (2∼100 dynes/cm^2^) (2); LPHN2 can convert force stimuli into membrane excitability by coupling to TMC1 and eliciting neurotransmitter glutamate release (3); and finally, specifically knocking out Lphn2 in hair cells causes hearing loss in mice (4). Therefore, our results suggest that GPCRs, such as LPHN2, could be auditory receptor candidates.

### Limitations of the Study

In this study, we have revealed an indispensable role of LPHN2 in auditory perception. However, it remains unclear whether LPHN2 mutations are correlated with auditory defects. Therefore, it will be important to revisit the potential pathogenetic roles of LPHN2 mutations in hearing loss through genome-wide association meta-analysis and exome sequencing. Despite our results indicating that the force applied on LPHN2 through LPHN-M-beads stimulates Ca^2+^ responses and glutamate release in hair cells *ex vivo*, further studies are warranted to investigate whether and how LPHN2 in cochlear hair cells converts the natural auditory stimuli into intracellular chemical signals and neurotransmitter release. Although the magnetic-bead-delivered force is theoretically within the physiological range, the relatively low binding efficiency of the beads with endogenous LPHN2, as well as the potential time delay in transmitting force, obscure its physiological significance. To this end, the force stimulus delivered by a fluid jet or stiff probe, which mimics the natural mechanical stimulation on hair cells, is required in future studies to determine the roles of LPHN2-mediated force sensation in Ca^2+^ response or neurotransmitter release. Finally, our current results suggest a physical and functional coupling between LPHN2 and TMC1. However, much remains unclear about the interaction mode between these two multi-transmembrane proteins as well as the potential relationship between the LPHN2-mediated TMC1 activation and tip link-mediated TMC1 activation. Further in-depth investigation using cryo-electron microscopy, or *in situ* cryo-electron tomography, combined with molecular dynamic simulations will shed light on the detailed structural information, which is expected to provide potential modulatory strategies for treating auditory defects by targeting LPHN2.

## Data availability

All data are available upon reasonable request from the corresponding authors.

## Acknowledgments

This work was supported by National Key R&D Program of China (2019YFA0904200 to J.-P.S., 2023YFA1801100 to Z.Y.), National Science Fund for Distinguished Young Scholars Grant (81825022 to J.-P.S., 82225011 to X.Y.), National Natural Science Foundation of China (32130055 and 82330118 to J.-P.S., 82330022 to X.Y., 81770407 to C.W., 82271176 to W.-W.L.), the Key Research Project of the Beijing Natural Science Foundation, China (Z200019 to J.-P.S.), Major Fundamental Research Program of Natural Science Foundation of Shandong Province, China (ZR2020ZD39 to J.-P.S. Z.-G.X, and W.-W.L, ZR2021ZD18 to X.Y.), Shandong Provincial Natural Science Fund (ZR2023QH394 to M.-W.W.), Taishan Scholars Program of Shandong Province (tsqn201909189 to W.-W.L.), Key Project of Precision Medicine Joint Fund of Natural Science Foundation of Hebei Province (81770407 H2020206409 to C.W.) and the Cutting Edge Development Fund of Advanced Medical Research Institute (to Z.Y., K.-K.Z. and J.-P.S.). J.-P.S. was also supported by the Tencent New Cornerstone Investigator Program.

## Author contributions

J.-P.S. and X.Y. initiated, designed and supervised the overall project. J.-P.S. conceived the idea of searching for MET-regulating GPCRs in the cochlear system at 2011 and initiated the screening process by collaborating with Z.-G.X.. X.Y., C.W. and W.X. designed and supervised all electrophysiology experiments. Z.-G.X and R.-J.C. provided guidance and suggestions on the cochlear studies. Z.Y. and J.-P.S. developed the magnetic force stimulation assay to examine the activation of G proteins and second messengers downstream of GPCRs. Z.Y., M.-W.W., S.-H.Z. and Z.-C.S. performed the cell-based assay of LPHNs. Z.Y., Z.-C.S., S.-H.Z., Q.-Y.Z., J.-R.G. and Y.-Q.W. performed the electrophysiological experiments supervised by J.-P.S. and X.Y.. Z.Y., M.-W.W., S.-H.Z., Z.-C.S., Q.-Y.Z. and X.-H.W. performed the Ca^2+^ imaging. J.-P.S., Z.Y., M.-W.W. and S.-H.Z. participated in the design of gene knockout mice and modified AAV-ie. Z.Y., M.-W.W., Z.-C.S., Q.-Y.Z., W.-W.L. and L.X. performed cochlear behaviour studies. Z.Y., M.-W.W., Z.-C.S. and X.-H.W isolated mouse cochlea and performed immunofluorescence studies and ex vivo experiments. M.-W.W., S.-H.Z and Z.-C.S. performed single cell qRT-PCR. M.-W.W. and Y. G. performed coimmunoprecipitation and western blotting analyses. Z.Y. and Y.G. performed BRET experiments. Z.Y., M.-W.W., S.-H.Z., Z.-C.S., J.-R.G. and W.-W.L. performed the AAV-ie-related experiments. K.-K.Z. and X.W. performed virtual screening and cellular assays under the supervision of J.-P.S. and X.Y.. J.-P.S., X.Y., Z.-G.X, C.W., R.-J.C, Z.Y., M.-W.W., S.-H.Z. and Z.-C.S. participated in data analysis and interpretation. Z.-G.X., C.W., W.X., W.-W.L. and L.X. provided insightful ideas related to the auditory sensation. Z.Y., M.-W.W., S.-H.Z., Z.-C.S., K-.K.Z, X.W., Q.-Y.Z., Y.G. and J.-R.G. prepared the figures. J.-P.S., X.Y., Z.Y., Z.-G.X and C.W. wrote the manuscript. All the authors have seen and commented on the manuscript.

## Methods

### Mice

C57BL/6J wild-type mice were obtained from Jackson Laboratory. *Lphn2^+/-^* (S-KO-15867), *Lphn2^fl/fl^* (S-CKO-17378), *Tmc1^-/-^* mice (S-KO-18952), *Gnas^fl/fl^* and LPHN2^mCherry^ mice with C57BL/6J background were purchased from Cyagen (Suzhou, China). *Pou4f3-CreER* mice with C57BL/6J background were generated by GemPharmatech (Nanjing, China). The *Pou4f3-CreER* line was generated by placing the *CreERT2* element downstream of the endogenous coding sequence of Pou4f3, which were separated by a P2A sequence. The *Pou4f3-CreER^+/-^ Lphn2^fl/fl^* and *Pou4f3-CreER^+/-^Gnas^fl/fl^*mice were generated by crossing *Pou4f3-CreER^+/-^* mice with *Lphn2^fl/fl^* mice and *Gnas^fl/fl^* mice, respectively. For activation of Cre recombinase in *Pou4f3-CreER^+/-^* mice, the mice were treated with tamoxifen (75 mg/kg) dissolved in corn oil through round window membrane (RWM) injection at P25 (left ear) and P26 (right ear) consecutively. For activation of Cre recombinase in *Pou4f3-CreER^+/-^* mouse embryos, the pregnant mice at 14 days post coitum (E14) was treated with 100 mg/kg tamoxifen supplemented with 37.5 mg/kg progesterone for 3 consecutive days through *i.p.* injection. All mice were housed at Shandong University Animals Care Facility, with a 12 hr light/12 hr dark cycle and ad libitum access to water. Both male and female mice were used and were randomly assigned to experimental groups. All mice care and experiments were reviewed and approved by Animal Use Committee of Shandong University Cheeloo College of Medicine.

### Cell lines

Human embryonic kidney 293 (HEK293) cells were obtained from American Type Culture Collection (ATCC, Manassas, VA, USA) and were cultured in DMEM supplemented with 10 % fetal bovine serum, penicillin (100 IU/ml), and streptomycin (100 mg/ml) in 5 % CO_2_ at 37 °C. Cell transfection was performed with Lipofectamine 2000 (Invitrogen) according to the manufacturer’s instructions.

### RNA extraction and Quantitative reverse-transcriptase PCR

Total RNA was extracted from the brain or cochlear epithelium of WT mice using TRIzol reagent (Invitrogen). cDNA was synthesized using the qRT-PCR Kit (Toyobo, FSQ-101) and the quantitative reverse-transcriptase PCR (qRT-PCR) was conducted using FastStart SYBR Green Master (Roche) on a LightCycler qPCR system (Bio-Rad). The relative mRNA levels of target genes, including *Lphn2*, *Tmc1* and *Gnas* subtypes and G protein subtypes, were calculated using *Actb* as an internal control. For single cell qRT-PCR, RNA was extracted from single cochlear hair cell (or AAV-ie-*Lphn2pr-*mCherry-labeled cochlear hair cell) using Discover-sc® WTA Kit V2 (Vazyme, N711). The cDNA synthesis, qRT-PCR and data processing were performed as described above. The sequences of all the primers used in the present study are listed in Table S1.

### Stereocilia separation

The stereocilia was separated from the hair cell body using a modified twist-off method as described previously (Avenarius et al., 2017; Gillespie and Hudspeth, 1991). Briefly, circular coverslip sheet with a diameter of 14 mm was soaked with a 4% ammonium hydroxide and 4% hydrogen peroxide solution, washed with water, and soaked in methanol. The coverslip was air-dried and then coated with 1 µL Corning® Cell-Tak™ Cell and Tissue Adhesive. Dissected basilar membrane of mouse cochlea was dropped onto the coverslips in dissection media and gently pressed to the coverslip with the epithelium side down. The basilar membrane was then removed, and stereocilia samples that are stuck on the coverslip were immunostained by phalloidin or gently scraped off for further analysis.

### Western blotting

The brain, cochlea or utricle from WT, *Lphn2^+/-^*, *Lphn2^-/-^*, *Pou4f3-CreER^+/-^Lphn2^+/+^* or *Pou4f3-CreER^+/-^Lphn2^fl/fl^* mice, or the isolated stereocilia and cell bodies of cochlear hair cells, or HEK293 cells transfected with LPHNs or empty plasmid pcDNA3.1 were collected with NP40 lysis buffer containing phosphatase and protease inhibitors. After centrifugation at 12,000 rpm for 20 min at 4 °C, the supernatant containing target proteins were collected. After determining the protein concentrations by BCA assay, the proteins were subjected to western blotting. Antibodies used in this study include anti-LPHN2 Rabbit antibody (Boster A30833, 1:1000), anti-VLGR1 Goat antibody (Santa Cruz sc-21252, 1:1000), anti-Myosin7a Mouse antibody (DHSB 138-1, 1:1000), anti-PCDH15 Rabbit antibody (Biomatik CAU22931, 1:1000), anti-CDH23 Mouse antibody (Santa cruz SC-166066, 1:1000), anti-FLAG Mouse antibody (Sigma F1804, 1:1000), anti-TMC1 Rabbit antibody (Abcam ab199949, 1:1000), anti-LHFPL5 Rabbit antibody (Thermo Fisher Scientific PA5-23919, 1:1000), anti-TMIE Rabbit antibody (Thermo Fisher Scientific PA5-58330, 1:1000), and anti-GAPDH Rabbit antibody (Abways AB0037, 1:10000).

### Immunofluorescence staining and confocal microscopy

For whole-mount immunostaining, the mouse temporal bones of WT, *Pou4f3-CreER^+/-^ Lphn2^+/+^*, *Pou4f3-CreER^+/-^Lphn2^fl/fl^*, or the *Pou4f3-CreER^+/-^Lphn2^fl/fl^* mice treated with AAV-ie-LPHN2 or AAV-ie-*Lphn2pr*-mCherry were fixed with 4% paraformaldehyde overnight at 4 °C. The cochlea was dissected as whole and treated with 0.5 M EDTA for 3∼5 h at 37 °C for decalcification. After three washing with PBS, the cochlea was permeabilized and blocked with PBS containing 0.5 % Triton-X and 5 % goat serum for 2 h. The cochlea whole mount was incubated with primary antibodies overnight at 4 °C, followed by incubation with secondary antibodies for 1∼2 h at 37 °C. The samples were washed three times with PBS, mounted in ProLong Gold Antifade (Thermo Fisher Scientific, Waltham, MA), and subjected to fluorescence microscopic analysis using a LSM980 laser scanning confocal microscope system (ZEISS, Germany). Image acquisition and merging were performed using ZEISS Zen software. For the cryosection immunostaining, the decalcified cochlea was sequentially dehydrated using sucrose solution with concentration gradient (15 %, 20 % and 30 % in PBS). The cochlea was embedded in Tissue-Tek OCT compound (Sakura Fintek USA, Torrance, CA) and 6 μm sections were cut. The cochlea slices were incubated with blocking buffer at 37 °C for 2 h and were processed for immunostaining and fluorescence microscopic analysis.

To investigate the subcellular localization of TMC1 in HEK293 cells, the cells were transiently transfected with plasmids encoding TMC1 and different GPCRs. 24 hours after transfection, the cells were seeded into 35-mm fibronectin-coated glass-bottom dish at the density of 5×10^5^ cells/dish. After another 24 hours, the cells were fixed and processed for immunostaining and fluorescence microscopic analysis.

Primary antibodies used in the present study were as follows: anti-LPHN2 Rabbit antibody (Sigma HPA043447, 1:200), anti-Myosin7a Mouse antibody (DHSB 138-1, 1:800), Phalloidin-iFluor 594 (Abcam ab176757, 1:5000), Phalloidin-iFluor 488 (Abcam ab176753, 1:5000), Phalloidin-iFluor 647 (Abcam ab176759, 1:5000), mouse anti-spectrin α (Sigma-Aldrich, ZRB2080, 1:100), anti-PCDH15 Rabbit antibody (Abcam ab202560, 1:100), anti-CDH23 Mouse antibody (Santa cruz SC-166066, 1:100), anti-TMC1 Rabbit antibody (Merck ABN1649, 1:100), anti-TMC2 Rabbit antibody (Abcam ab200039, 1:100), anti-LHFPL5 Rabbit antibody (Thermo Fisher Scientific PA5-23919, 1:100), anti-TMIE Rabbit antibody (Thermo Fisher Scientific PA5-58330, 1:100), and anti-FLAG Mouse antibody (Sigma F1804, 1:100). Secondary antibodies were as follows: Alexa Fluor Plus 555 Goat anti-Mouse IgG (Invitrogen, A32727), Alexa Fluor Plus 488 Goat anti-Rabbit IgG (Invitrogen, A32731) and Alexa Fluor Plus 647 Goat anti-Mouse IgG (Invitrogen, A32728).

### Auditory brainstem response (ABR) measurements

The *Lphn2^+/-^*, *Pou4f3-CreER^+/-^Lphn2^fl/fl^*mice or their wild-type littermates at P30, or the *Pou4f3-CreER^+/-^Lphn2^fl/fl^* mice treated with AAV-ie-LPHN2 or control virus at P30 were placed on an isothermal pad to keep the body temperature at 37 °C during the experiment. After anesthetization with ketamine (100 mg/kg) and xylazine (25 mg/kg) intraperitoneally, electrodes were inserted subcutaneously at the vertex and pinna as well as near the tail. The stimulus generation, presentation, ABR acquisition, and data management were coordinated using an RZ6 workstation and BioSig software (Tucker Davis Technologies, Inc.). Acoustic stimuli (clicks or pure-tone bursts) of various sound levels were generated using high-frequency transducers. At each sound level, 512 responses were sampled and averaged. ABR thresholds were determined for each animal as the lowest sound level at which all ABR waves were detectable.

### Distortion product otoacoustic emission (DPOAE) measurements

Mice were anesthetized and maintained as described for ABR experiment. Two sine wave tones (f2 = 1.2 × f1) were presented for 1 sec durations at various sound levels. The emitted acoustic signal was then picked up by a microphone and digitized, and the magnitude of the distortion product (2 × f1-f2) was determined. The surrounding noise floor was calculated by averaging adjacent frequency bins around the frequency of the distortion product. DPOAE thresholds were determined as the lowest stimulus sound level at which the emitted signal of the distortion product was at least 5 dB SPL above the noise floor.

### Scanning Electron Microscopy (SEM)

Mouse temporal bones were fixed with 2.5 % glutaraldehyde in 0.1 M phosphate buffer overnight at 4 °C. Cochlea were dissected out of the temporal bone and post-fixed with 1 % osmium tetroxide in 0.1 M phosphate buffer at 4 °C for 2 h. Samples were then dehydrated in ethanol and critically point-dried using a Leica EM CPD300 (Leica, Germany). Samples were then mounted and sputter coated with platinum (15 nm) using a Cressington 108 sputter coater (Cressington, United Kingdom). The images were taken with a Quanta250 field-emission scanning electron microscope (FEI, The Netherlands).

### Endogenous cAMP measurement

To measure the cAMP levels in mouse cochlea or utricle or in HEK293 cells, the isolated cochlear or utricle explants or HEK293 cells transfected with Flag-LPHN2 were treated without or with LPHN2-M-beads for 15 min in the presence of IBMX. The harvested cochlear or utricle explants or HEK293 cells were homogenized in cold 0.1 M HCl and centrifuged at 12000 rpm for 10 min to remove the particulates. The supernatants were collected and neutralized with 1M NaOH. The cAMP levels were determined using cAMP ELISA Kit (R&D systems, KGE012B) according to manufacturer’s instructions.

### Structural modelling of LPHN2 and virtual screening for LPHN2 inhibitors

Homology modelling of LPHN2 was performed with the online program SWISS-MODEL (https://swissmodel.expasy.org/)(Waterhouse et al., 2018). First, the primary amino acid sequence of human LPHN2 (E835-C1098) was uploaded to the program, then the inactive CD97 structure was also uploaded as template for homology modelling of LPHN2 structure (Mao et al., 2024b). Model was built based on the target-template alignment using ProMod3. Coordinates which are conserved between the target and the template are copied from the template to the model. Insertions and deletions are remodeled using a fragment library. Side chains are then rebuilt. Finally, the geometry of the resulting model is regularized by using a force field. The global and per-residue model quality has been assessed using the QMEAN scoring function (Studer et al., 2020). The interactions between LPHN2 (I840-G1135) and TMC1 (C161-Q760) was simulated using the online program ColabFold v1.5.5 (AlphaFold2 using MMseqs2) (Mirdita et al., 2022).

The predicted ligand binding pocket in modelled structure of LPHN2 was analyzed through *SiteMap* algorithm with Maestro (version 13.4, Schrödinger LLC, New York, NY) software (Halgren, 2009). All atoms in modelled LPHN2 structures were considered as the receptor. The grid spacing was set to 0.7 Å, and points that border on too many assigned philic points were excluded. Each pocket required at least 15 site points, and top four potential binding pockets were identified and analyzed. Maestro software was employed to perform virtual screening. The modelled structure of LPHN2 was used to construct the virtual screening model. First, the Protein Preparation Workflow was used to prepare the modelled LPHN2 structure with default settings. Then, the docking grid of the predicted top 1 binding pocket was generated by the Receptor Grid Generation panel with default settings. Subsequently, the 1000,000 molecules in our designed compound database were prepared using the LigPrep panel (LigPrep, version 2.3) in Maestro with the following settings: the ionized states and tautomers were generated by Epik algorithm at pH 7.0 ± 2.0; The chiralities were retained for ligands with specified chiralities; The maximum number of stereoisomers for each ligand was set to 32. The OPLS4 force field was used for energy minimization. Finally, molecular docking was performed using the Ligand Docking panel with XP (extra precision) mode. In this process, all the parameters were set as the default settings. Clustering analysis was performed using “clustering Molecules” protocols integrated in Pipeline Pilot, version 7.5 (Pipeline Pilot; Accelrys SoftwareInc., San Diego, CA).

### BRET assay

#### G protein dissociation BRET assay

To investigate the force-induced G protein activation of LPHN2, the G protein dissociation BRET assay was performed as previously described (Yang et al., 2021; Yang et al., 2024). Briefly, HEK293 cells co-transfected with either LPHN2 or its mutations, along with Gs or Gi BRET probes were incubated for 24 hours. The cells were then plated into a 96-well microplate at a density of 5 × 10^5^ cells per well. The cells were incubated for another 24 hours at 37°C before being subjected to mechanical stimulation with a force of 3pN for 3 minutes. Following the mechanical stimulation, coelenterazine 400a (Interchim Cayman) at a final concentration of 5 μM was added, and BRET signals were measured using a Mithras LB940 multimode reader (Berthold Technologies). The BRET signal was determined as the ratio of light emission at 510 nm to that at 400 nm.

#### LPHN2-TMC1 BRET assay

To detect the constitutive interaction between LPHN2 and TMC1, HEK293 cells were transfected with a fixed amount of Myc-VLGR1a and LPHN2-Nluc (LPHN2-ICL1-Nluc, LPHN2-ICL2-Nluc, LPHN2-ICL3-Nluc or LPHN2-Cter-Nluc) along with an increasing concentration of TMC1-YFP (N ter-YFP-TMC1 or TMC1-Cter-YFP) and incubated for 48 h at 37 °C. Then the cells were digested, centrifuged and resuspended in BRET buffer (140 mM NaCl, 2.7 mM KCl, 1 mM CaCl_2_, 0.9 mM MgCl_2_, 0.37 mM NaH_2_PO_4_, 12 mM NaHCO_3_, 25 mM HEPES and 5.5 mM d-glucose), and then distributed into black-wall clear-bottom 96-well plates at a density of 5 ×10^4^ cells per well. After adding coelenterazine H (a final concentration of 5 mM), the light emissions at 540 nm and 460 nm were recorded using a Mithras LB 940 Multimode Microplate Reader. The BRET signal was calculated as the ratio of the Em540 over Em460. The net BRET was calculated by subtracting the BRET signal obtained in the cells transfected without TMC1-YFP.

For the force-induced interaction between LPHN2 and TMC1, the HEK293 cells transfected with Flag-LPHN2-Nluc, TMC1-YFP and VLGR1a were incubated with buffer containing polylysine-coated Ctrl-beads or Flag-M-beads for 15 min. The cells were stimulated with or without force for 2 min, and then coelenterazine H (a final concentration of 5 μM) was added, followed by the recording of light emissions at 540 nm and 460 nm as described above. The △BRET was calculated as the BRET signal change due to the force stimulation.

#### FlAsH-BRET assay

To investigate the force-induced conformational changes in TMC1, HEK293 cells were transfected with Myc-VLGR1a, Flag-LPHN2 and Nluc-TMC1-FlAsH probes (N-Nluc-TMC1-ICL1-FlAsH, N-Nluc-TMC1-ICL2-FlAsH, N-Nluc-TMC1-ICL3-FlAsH, or N-Nluc-TMC1-ICL4-FlAsH) and incubated for 24 h at 37 °C. Then the cells were distributed into 96-well plates at a density of 5 ×10^4^ cells per well. After 24 h, the cells were labeled with FlAsH-EDT2 (a final concentration of 2.5 μM) for 40 min at 37 °C. After washing twice with PBS, the cells were treated with polylysine-coated Ctrl-beads or Flag-M-beads for 15 min in the absence or presence of NF449 (10 μM) or D11 (50 nM). The cells were then stimulated with magnetic force for 2 min. After adding coelenterazine H (a final concentration of 5 μM), the light emissions at 540 nm and 460 nm were recorded. The BRET signal was calculated as the ratio of the Em540 over Em460.

### Cell surface ELISA assay

The cell surface expression levels of LPHN2 and its mutants, or TMC1 conformational sensors were determined by cell surface ELISA assay. Briefly, HEK293 cells transiently transfected with N-terminal Flag-tagged wild-type LPHN2 or mutants, or TMC1 conformational sensors in 24-well plates for 48 h were fixed with 4% (w/v) paraformaldehyde and incubated with anti-Flag M2 primary antibody and a secondary goat anti-mouse antibody conjugated to horseradish peroxide. A color reaction was initiated by adding TMB (3,3′,5,5′-Tetramethylbenzidine) solution and stopped by adding an equal volume of 0.25 M HCl. The absorbance at 450 nm of each well was measured to determine the relative expression levels. The transfecting amounts of the plasmids encoding corresponding mutants or conformational sensors were adjusted to enable equal cell-surface expression levels of the target proteins.

### Adeno-associated virus preparation and injection

The AAV-ie-*Lphn2pr*-mCherry (3.5 × 10^13^ genomic copies per ml) and AAV-ie-LPHN2 (3.0 × 10^13^ genomic copies per ml)) were generated by OBiO Technology (Shanghai, China). Both viruses were modified from the original AAV-ie vectors (Tan et al., 2019). Briefly, for the construction of AAV-ie-*Lphn2pr*-mCherry, the original CAG promoter in the AAV-ie was replaced with the promoter region (800 bp) of *Lphn2*, which directs the expression of mCherry and could be used for the labelling of LPHN2-expressing cochlear hair cells. For in vivo rescue experiment, the LPHN2-GAIN sequence with an N-terminal 3×Flag tag was inserted into the AAV-ie-*Lphn2pr*-mCherry vector. An IRES element was placed between the Flag-LPHN2-GAIN and the mCherry coding sequence. All viral vectors were aliquoted and stored at -80 °C until use.

For virus injection, *Pou4f3-CreER^+/-^Lphn2^+/+^*or *Pou4f3-CreER^+/-^Lphn2^fl/fl^* mice at P3 were placed in an ice bath until loss of consciousness and then moved to an ice pad for surgical procedures. Injections were performed through the round window membrane of left ear of mouse with a glass micropipette (25 μm) at a speed of 1 μL virus / min controlled by a micromanipulator UMP3 UltraMicroPump (World Precision Instruments). A total volume of 0.8 μL of the AAV-ie-*Lphn2pr*-mCherry or AAV-ie-LPHN2 virus was injected and then the incision was closed using veterinary tissue adhesive (Millpledge Ltd, UK). The mice were placed on a 37 °C warming pad until fully awake.

### Single-cell Ca^2+^ imaging

The cochlear explants derived from *Pou4f3-CreER^+/-^Lphn2^+/+^*or *Pou4f3-CreER^+/-^Lphn2^fl/fl^* mice were attached to 12-mm round glass coverslips, which were pre-coated with polylysine and placed in 35-mm dishes. Cochlear explants were incubated with imaging buffer (10 mM glucose,150 mM NaCl, 5 mM KCl, 1.3 mM MgCl_2_, 1.2 mM NaH_2_PO_4_, 3 mM CaCl_2_, 20 mM HEPES, pH 7.4) containing 2.5 μM Fura-2 and 0.05 % Pluronic F-127 (Life Technologies) for 15 min at 37 °C. After washing with imaging buffer, the cochlear explants were incubated with fresh imaging buffer containing LPHN2-M-beads or Ctrl-beads for 20 min at 37 °C to enable the association of the antibody-coated beads with the target proteins. Then, the dish containing the cochlear explants was mounted into an Olympus IX71 microscope equipped with a FluoCa BioHD CMOS camera and a pE-340fura illumination system (coolLED). The cochlear hair cells labelled by AAV-ie-*Lphn2pr*-mCherry were selected for Ca^2+^ imaging. The F340/F380 ratio of selected cells were recorded while the magnetic force was applied on the LPHN2-M-beads or Ctrl-beads. Data are recorded and analyzed using the VisiFLUOR Fluorescence Ratio Imaging System (Visitron Systems).

### Glutamate secretion detection

To characterize the glutamate secretion from cochlear hair cells, AAV-ie-*hSyn*-R^ncp^-iGluSnFR (4 × 10^13^ genomic copies per ml) were delivered to *Pou4f3-CreER^+/-^Lphn2^+/+^*or *Pou4f3-CreER^+/-^Lphn2^fl/fl^* mice at P3. A total volume of 0.8 μL of the AAV-ie-*hSyn*-R^ncp^-iGluSnFR virus was injected and then the incision was closed using veterinary tissue adhesive (Millpledge Ltd, UK). The cochlea was isolated at P10 and the explants were incubated with fresh imaging buffer containing LPHN2-M-beads or Ctrl-beads for 20 min at 37 °C to enable the association of the antibody-coated beads with the target proteins. Widefield imaging was performed on an inverted Olympus IX71 microscope equipped with a 200 W metal halide lamp (PRIOR Lumen), 60× objectives, and a 16-bit QuantEM 512SC electron-multiplying CCD camera (Photometrics). A filter set of 620/60 nm (emission) and 570 nm (dichroic) was used for recording cells labelled by AAV-ie-*hSyn*-R^ncp^-iGluSnFR. The selected cells were recorded while the magnetic force was applied on the LPHN2-M-beads or Ctrl-beads. Data are recorded and analyzed using the VisiFLUOR Fluorescence Ratio Imaging System (Visitron Systems).

### Electrophysiology

AAV-ie-*Lphn2pr*-mCherry was delivered into *Pou4f3-CreER^+/-^Lphn2^+/+^* or *Pou4f3-CreER^+/-^Lphn2^fl/fl^* mice at P3. The organs of Corti were isolated at P10 and the mCherry-labelled OHCs and IHCs were selected for electrophysiological studies. Excised cochlear turns were placed in a recording chamber mounted on an upright microscope (Olympus), and visualized using a 63 x 0.9 NA water-immersion objective and infrared differential interference contrast. MET currents were recorded using fluid jet assay as described previously (Liu et al., 2019). Briefly, currents were evoked using a fluid jet from a pipette (tip diameter of 5–10 μm) positioned facing the staircase side of the hair bundle at a distance of around 5 μm at room temperature. The fluid jet was driven by a 40-Hz sinusoidal voltage to deflect the hair bundles. The fluid flow was controlled by a 27-mm-diameter piezoelectric disk driven by a homemade 20 × amplifier that precisely outputs analog driving voltage. The command signal was generated by an amplifier (EPC10, HEKA) controlled by the PatchMaster software with a holding potential of –80mV. Patch pipettes were pulled from borosilicate glass (ITEM #:BF150-86-10, Sutter Instrument Co.) using a micropipette puller (P-97, Sutter Instrument Co.) to a resistance of 3-5 MΩ. The pipette solution contained 142 mM KCl, 3.5 mM MgCl_2_, 1 mM EGTA, 2.5 mM MgATP, 0.1 mM CaCl_2_, 5 mM HEPES, adjusted to a pH of 7.4 with KOH. The external solution contained 145 mM NaCl, 5.8 mM KCl, 0.9 mM MgCl_2_, 1.3 mM CaCl_2_, 0.7 mM NaH_2_PO_4_, 10 mM HEPES and 5.6 mM D-glucose, adjusted to a pH of 7.4 with NaOH.

For the electrophysiology in the heterologous system, HEK293 cells transfected with GPCR-HC1, TMC1 and LPHN2 were plated onto coverslips coated with polylysine. The single channel currents in HEK293 cells were recorded 24-48 h after transfection. Cell-attached patch recordings were performed in extracellular solution (140 mM NaCl, 5 mM KCl, 2.5 mM CaCl_2_, 1 mM MgCl_2_, 10 mM glucose, 10 mM HEPES, pH 7.4 with NaOH) using 6-8 MΩ resistance electrodes filled with intracellular solution (140 mM CsCl, 5 mM EGTA and 10 mM HEPES, pH 7.4 with KOH). The current-voltage (I-V) curves of the spontaneous currents were plotted and fitted to the response amplitude measured from -80 to +80 mV in 20-mV increments. The single-channel conductance was obtained through linear fitting of the current-voltage plots. The inhibitors were added to the bath solution and the channel opening was recorded again from the same patch. The following inhibitors were tested: ruthenium red (RR, 40 μM), FM1-43 (3 μM), dihydrostreptomycin (DHS, 0.2 mM), amiloride (0.2 mM), NF449 (20 μM) and D11 (50 nM). For the force-induced single-channel currents, the cells were incubated with fresh extracellular solution containing LPHN2-M-beads or CDH23-M-beads for 20 min at 37 °C to enable the association of the antibody-coated beads with the target proteins. Then, the cell-attached patch recordings were performed while the magnetic force was applied on the LPHN2-M-beads or Ctrl-beads.

### Co-immunoprecipitation

The cochlea from WT mice were ground and lysed with NP40 lysis buffer containing phosphatase and protease inhibitors. The lysates were immunoprecipitated using LPHN2- or control-antibody-conjugated agarose overnight at 4 °C. After washing with PBS, the immunoprecipitated proteins were subjected to electrophoresis and western blotting analysis using specific antibodies against specific MET channel components, including TMC1, PCDH15, LHFPL5, and TMIE.

### Mass spectrometry analysis

The cochlear epithelium derived from wild-type mice were homogenized using NP40 lysis buffer, supplemented with phosphatase and protease inhibitors. The resulting lysates were subjected to immunoprecipitation for 4 hours at 4°C using LPHN2-GAIN-bait (LPHN2-GAIN conjugated to the anti-Flag M2 beads) or control bait (empty anti-Flag M2 beads). After a 30 min treatment with 10% DSP on a shaker, the cross-linking reaction was terminated by the addition of 1 M Tris (pH 7.5). Following centrifugation at 1000 rpm for 5 min at 4°C, the precipitated beads were collected and subjected to three washes with a wash buffer (20 mM HEPES pH 7.5, 4 mM MgCl_2_, 15% Glycerin, 3 mM CaCl_2_, 100 mM NaCl, 0.01% MNG, 0.006% CHS) containing phosphatase and protease inhibitors. The precipitated beads were then eluted using the wash buffer containing 10 mM EGTA and 0.1 mg/ml Flag peptide, and subsequently concentrated using a concentrating column. Finally, the samples were stained using a silver staining kit and digested with trypsin for subsequent mass spectrometry analysis (Shanghai Applied Protein Technology Co., Ltd).

### Statistical analysis

All data in the present study were presented as mean ± SEM values with the numbers of mice or individual experiments indicted in the Figure or Figure legends. Unpaired two-tailed Student’s *t* test was used for the comparison between two groups. Differences between multiple groups with one or two variables were determined using one-way or two-way ANOVA followed by Dunnett’s post hoc test, respectively. Statistical analysis was performed using GraphPad Prism 9 software. Significance was defined as *P < 0.05, **P < 0.01, ***P < 0.001. No methods were used to determine whether the data met assumptions of the statistical approach. No power analysis was performed to determine the sample size. The sample size of mice in each experiment was determined based on experience from previous studies in our lab.

